# Dose-dependent nuclear delivery and transcriptional repression with a cell-penetrant MeCP2

**DOI:** 10.1101/2022.06.03.494754

**Authors:** Xizi Zhang, Madeline Zoltek, Deepto Mozumdar, Alanna Schepartz

**Affiliations:** Department of Chemistry, University of California, Berkeley, CA 94720; Department of Molecular and Cellular Biology, University of California, Berkeley, CA 94720; Department of Chemistry, Yale University, New Haven, CT 06520; California Institute for Quantitative Biosciences (QB3), University of California, Berkeley, CA 94720; Chan Zuckerberg Biohub, San Francisco, CA 94158

**Author notes:** **Materials & Correspondence.** Correspondence to Alanna Schepartz.

## Abstract

Methyl-CpG-binding-protein 2 (MeCP2) is a nuclear protein expressed in all cell types, especially neurons^1^. Mutations in the *MECP2* gene cause Rett syndrome (RTT), an incurable neurological disorder that disproportionately affects young girls^2^. Strategies to restore MeCP2 expression phenotypically reverse RTT-like symptoms in male and female MeCP2-deficient mice^3–5^, suggesting that direct nuclear delivery of functional MeCP2 could restore MeCP2 activity. We report that ZF-*t*MeCP2, a conjugate of MeCP2(Δaa13-71, 313-484)^6^ and the cell-permeant mini-protein ZF5.3^7–11^, both binds DNA in a methylation-dependent manner and reaches the nucleus of model cell lines intact at concentrations above 700 nM. When delivered to live cells, ZF-*t*MeCP2 engages the NCoR/SMRT co-repressor complex and selectively represses transcription from methylated promoters. Efficient nuclear delivery of ZF-*t*MeCP2 relies on a unique endosomal escape portal provided by HOPS-dependent endosomal fusion. The Tat conjugate of MeCP2 (Tat-*t*MeCP2), evaluated for comparison, is degraded within the nucleus, is not selective for methylated promoters, and trafficks in a HOPS-independent manner. These results support the feasibility of a HOPS-dependent portal for delivering functional macromolecules to the cell interior using the cell-penetrant mini-protein ZF5.3. Such a strategy could broaden the impact of multiple families of protein-derived therapeutics.

## Introduction

Methyl-CpG-binding-protein 2 (MeCP2) is an abundant nuclear protein expressed in all cell types, especially neurons^1^. Mutations in the *MECP2* gene cause Rett syndrome (RTT), a severe and incurable neurological disorder that disproportionately affects young girls^2^. Many potential RTT treatments are under development^12^, but no disease modifying treatment yet exists. Two features of RTT etiology render therapeutic development especially challenging. The first is that more than 850 different mutations in the *MECP2* gene^13^ account for > 95% of classical RTT cases^14^; this feature complicates approaches based on gene-editing^15–17^. The second is that excess MeCP2 protein causes MeCP2 duplication syndrome^18^ which itself causes progressive neurological disorders; this feature complicates approaches that rely on gene delivery^19,6,20–22^. Since 2007, several studies have demonstrated that restoring MeCP2 expression can phenotypically reverse RTT-like symptoms in male and female MeCP2-deficient mice^3–5^. These rescue experiments provide evidence that dose-dependent, nuclear delivery of functional MeCP2 protein could provide a novel treatment modality. Although the concentration of MeCP2 varies between cell types, its primary function is to engage the NCoR/SMRT co-repressor complex in a methylated DNA-dependent manner^23^. Thus, in order to be effective, MeCP2 protein must reach the nucleus intact, transcriptionally active, and in the high nanomolar to low micromolar concentration range^1,24^.

Previous efforts to deliver MeCP2 protein suffered from low delivery efficiency and significant cargo degradation^25–29^. We showed previously that the mini-protein ZF5.3^7–11^ is taken up by the endosomal pathway and released efficiently into the cytosol and nuclei of live cells^8^, alone and when fused to certain protein cargos. Proteins successfully delivered using ZF5.3 include the model protein SNAP-tag^9^, the metabolic enzyme argininosuccinate synthetase^11^, and the proximity labeling tool APEX2^9^. These proteins differ in molecular weight, stoichiometry, isoelectric point, and the presence of bound cofactors. In all cases evaluated, the protein that reached the cytosol was fully intact as judged by Western blot analysis of isolated cytosolic fractions free of detectable endosomal contamination, and the delivery efficiencies were 2-10-fold^9^ higher than seen with canonical or cyclic peptides^30^. Mechanistic studies confirm that ZF5.3 relies on the endocytic pathway to reach the cell interior^7^, and that endosomal escape into the cytosol demands a functional homotypic fusion and protein sorting (HOPS) complex^10^.

Here we use chemical biology, cell biology, biophysics, and biochemistry tools to qualitatively and quantitatively assess the nuclear delivery and function of MeCP2 conjugates of ZF5.3 and Tat. Although both conjugates bind DNA in a methylation-dependent manner *in vitro* and appear to reach the nucleus as judged by fluorescence-based methods, biochemical fractionation studies reveal that only the conjugate with ZF5.3 remains fully intact within the nucleus. When delivered to live cells, the conjugate between ZF5.3 and MeCP2 effectively engages the NCoR/SMRT co-repressor complex and selectively represses transcription from methylated promoters. Efficient nuclear delivery relies on HOPS-dependent endosomal fusion. By contrast, the Tat conjugate of MeCP2 is degraded within the nucleus, is not selective for methylated promoters, and trafficks in a HOPS-independent manner. The results described here support the feasibility of a HOPS-dependent portal for delivering functional macromolecules to the cell interior using the cell-penetrant mini-protein ZF5.3. Such a strategy could broaden the impact of multiple families of protein-derived therapeutics.

## Results

### Design, purification, and characterization of MeCP2 variants

Full-length murine MeCP2 (MeCP2-e2) contains 484 amino acids (52 kD) (**Fig. 1a**)^6^. MeCP2(ΔNC) (referred to henceforth as *t*MeCP2) is shorter (27 kDa, 253 aa) but mirrors MeCP2 in its interactions with methylated DNA and the NCoR/SMRT complex, and *Mecp2*-null male mice display a near-normal phenotype upon expression of MeCP2(ΔNC)^6^. We generated fusion proteins containing a single copy of ZF5.3^7^ or Tat_47-57_ ^31^ followed by the complete sequence of *t*MeCP2 (**Fig. 1a**). Each fusion protein also contained a sortase recognition motif (6 aa) to enable site-specific fluorophore conjugation and a Strep-tag II sequence (8 aa) to enable affinity purification. We also prepared two *t*MeCP2 variants with substitutions that alter function. The first is T158M *t*MeCP2, with a methyl-CpG-binding domain (MBD) mutation that reduces specific DNA binding^32,33^ and is seen commonly in RTT patients^14^. The second is P302L *t*MeCP2, which has a diminished ability to engage the TBLR1 subunit of the NCoR/SMRT repressor complex^34^. All *t*MeCP2 variants were expressed in *E. coli*, purified to > 95% homogeneity, and characterized using Western blots and LC/MS (**Extended Data Fig. 1a, b**). Variants carrying a fluorescent label were generated using sortase-A and a GGGK-lissamine rhodamine B (Rho) co-reagent as previously described^9^ (**Extended Data Fig. 1a, b**). Circular dichroism (CD) analysis of all *t*MeCP2 variants confirmed that the conjugation of ZF5.3 and Tat_47-57_ had minimal effect on protein secondary structure (**Extended Data Fig. 1c)**. Consistent with previous reports for full-length MeCP2^35^, all *t*MeCP2 variants show high levels of intrinsic disorder (60%) in the absence of DNA (**Supplementary Table 1**).

**Fig. 1.**
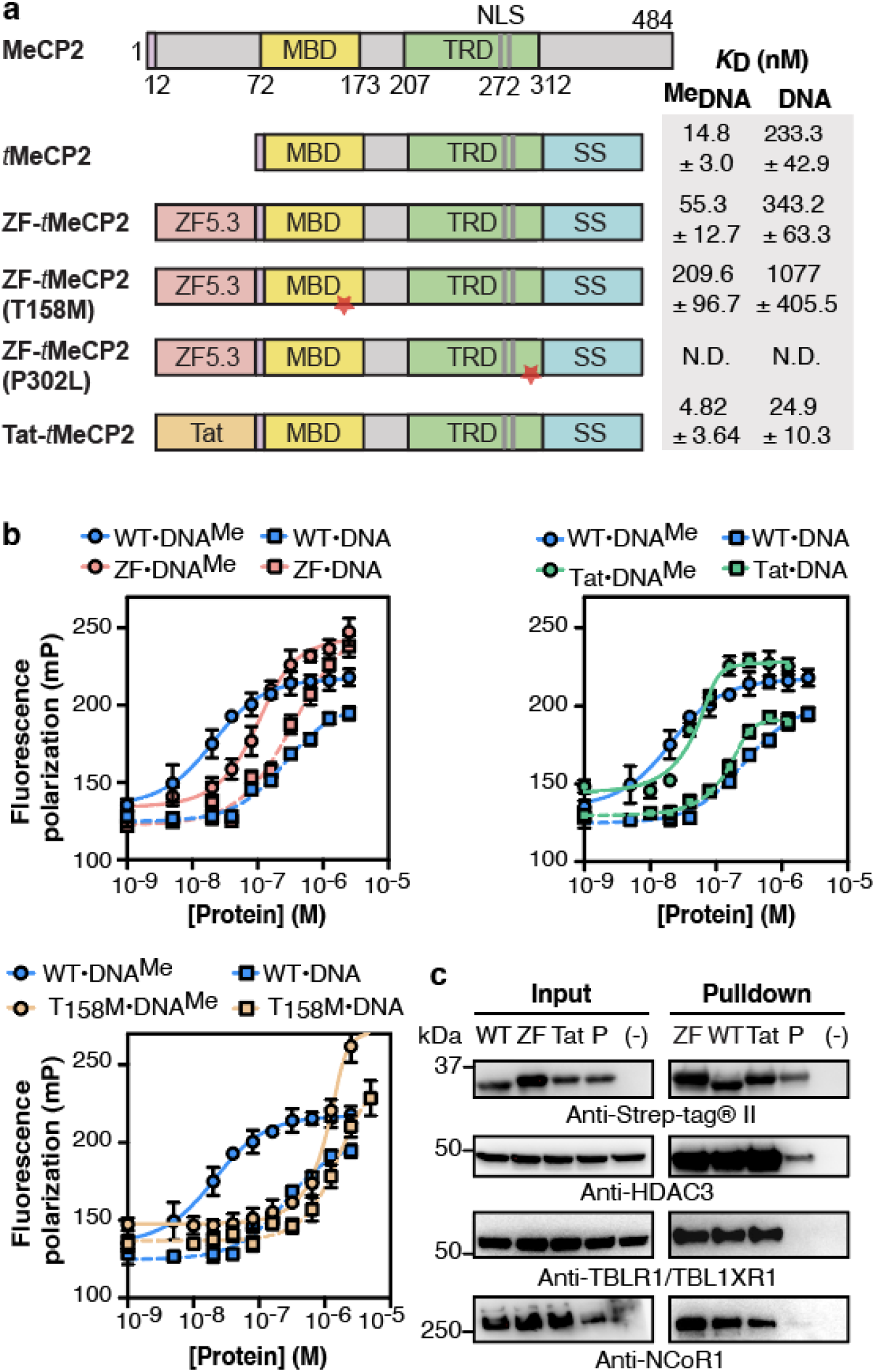
ZF-*t*MeCP2 is functional *in vitro*. **a**, *t*MeCP2 proteins used in this work lack N-terminal residues 13-71 and C-terminal residues 313-484. All proteins were expressed in *E. coli* and purified as described in Methods and Materials. MBD: methyl-CpG-binding domain; TRD: transcriptional-repressor domain; NLS: nuclear localization sequence; SS: sortase motif + Strep-tag. The red star indicates the location of the point mutation. **b**, Plots showing changes in fluorescence polarization used to calculate the apparent equilibrium dissociation constant (*K*_D_) of the complex between each *t*MeCP2 variant and methylated (DNA^Me^) or non-methylated (DNA) oligonucleotides (**Supplementary Table 3**). The data were fitted to an equilibrium binding equation based on the Langmuir model^39^ to calculate the *K*_D_ values in **a (**error represents standard error). Data are represented as mean ± SD. n = 6 (three technical replicates each from two biological replicates). **c**, Western blots were used to analyze an *in vitro* anti-Strep-tag pull-down assay to probe the interaction of the indicated *t*MeCP2 variant (WT: *t*MeCP2, ZF: ZF-*t*MeCP2, Tat: Tat-*t*MeCP2, P: ZF-*t*MeCP2(P302L)) with subunits of the NCoR/SMRT repressor complex. HDAC3: Histone Deacetylase 3; TBLXR1: Transducin Beta-Like 1X-Related Protein 1; NCoR1: Nuclear Receptor Corepressor 1. The gel results shown are representative of three biological replicates.

### Purified *t*MeCP2 proteins are active *in vitro*

MeCP2 functions like a bridge to repress transcription from methylated promoters^23^. The N-terminal methyl-CpG-binding domain (MBD) interacts with methylated DNA^32^ while the C-terminal transcriptional-repressor domain (TRD) domain engages NCoR1/2 co-repressor partners^34,36^. To establish whether the *t*MeCP2 proteins studied here retain these functions *in vitro*, we measured their affinities for methylated and non-methylated DNA oligonucleotides using a fluorescence polarization assay and used immunoprecipitation methods to assess interactions with co-receptor proteins in lysates (**Fig. 1b, c**). Fluorescence polarization analysis was performed with a 22 bp double-stranded, fluorescein-tagged, DNA oligonucleotide containing a methylated or non-methylated cytosine (**Fig. 1b**). *t*MeCP2 interacts with methylated DNA with a *K*_D_ of 15 nM and a 15-fold preference for methylated versus non-methylated DNA. ZF-*t*MeCP2 interacts with methylated DNA with a *K*_D_ of 55 nM and a 6-fold preference for methylated DNA. The conjugate of Tat_47-57_ and *t*MeCP2 (Tat-*t*MeCP2) interacts with both methylated DNA (*K*_D_ = 5 nM) and non-methylated DNA (*K*_D_ = 25 nM) more favorably than *t*MeCP2 and ZF-*t*MeCP2 with a 5-fold preference for methylated DNA. As expected, ZF-*t*MeCP2(T158M) binds poorly to both methylated (*K*_D_ = 210 nM) and non-methylated (*K*_D_ = 1.1 μM) DNA when compared to *t*MeCP2. Although the *K*_D_ describing the interaction of *t*MeCP2 with methylated DNA has not previously been determined, reported values for full-length MeCP2 fall in the range of 36-130 nM with a 2-33 fold preference for methylated DNA^37,38^.

To further probe the function of purified ZF-*t*MeCP2 *in vitro*, we used affinity pull-down assays to evaluate its interactions with the NCoR/SMRT co-repressor complex in nuclear lysates of NIH3T3 cells^6^. Lysates^36^ were incubated overnight at 4 °C with 1.5 μM of ZF-*t*MeCP2, Tat-*t*MeCP2, or *t*MeCP2; ZF-*t*MeCP2(P302L) was used as a negative control. Each *t*MeCP2 variant was extracted from the lysates using streptavidin-coated beads, and the identities and relative levels of bound NCoR/SMRT subunits (NCoR1, HDAC3, and TBL1/TBLR1) were evaluated using Western blots (**Fig. 1c**). These blots revealed that *t*MeCP2, ZF-*t*MeCP2, and Tat-*t*MeCP2 remain intact after an overnight incubation with lysates at 4 °C and effectively sequester HDAC3, TBL1/TBLR1, NCoR1 from NIH3T3 nuclear cell lysates. In all cases, there was little or no evidence of interaction with ZF-*t*MeCP2(P302L). Taken together, these two *in vitro* assays confirm that purified ZF-*t*MeCP2 retains the core functions of MeCP2: selective recognition of methylated DNA and the ability to engage the NCoR/SMRT co-repressor complex.

### Efficient delivery of ZF-*t*MeCP2 to the nuclei of Saos-2 and CHO-K1 cells

Next, we made use of three fluorescence-based methods and two model cell lines to evaluate the overall uptake of each *t*MeCP2 variant and specifically how much protein trafficked to the nucleus, the site of MeCP2 function. Human osteosarcoma (Saos-2) cells were incubated for 1 hr with rhodamine-tagged *t*MeCP2-Rho, ZF-*t*MeCP2-Rho, Tat-*t*MeCP2-Rho, or ZF-*t*MeCP2(T158M)-Rho at concentrations between 0.5 μM and 2 μM (**Fig. 2a**). When visualized using 2D confocal microscopy, cells treated individually with each of the four *t*MeCP2-Rho variants showed bright punctate intracellular fluorescence, while no fluorescence was observed in non-treated cells (**Extended Data Fig. 2a**). Saos-2 cells treated with ZF-*t*MeCP2-Rho, Tat-*t*MeCP2-Rho, and ZF-*t*MeCP2(T158M)-Rho also showed evidence of intra-nuclear fluorescence at concentrations as low as 1 μM, while cells treated *t*MeCP2-Rho did not, even at 2 μM concentration. When visualized as 3D z-stacks, cells treated with ZF-*t*MeCP2-Rho, Tat-*t*MeCP2-Rho, and ZF-*t*MeCP2(T158M)-Rho differed in intra-nuclear localization (**Fig. 2b, Extended Data Fig. 2b, Supplementary Video 1-4**). Cells treated with ZF-*t*MeCP2-Rho and Tat-*t*MeCP2-Rho showed an even distribution of rhodamine fluorescence in Hoechst-positive, DNA-rich regions, whereas cells treated with ZF-*t*MeCP2(T158M)-Rho showed aggregated rhodamine signal in small discrete regions resembling nucleoli. This observation aligns with previous reports that truncation of the entire MBD or T158M mutation resulted in MeCP2 relocalization to the nucleolus^40,41^.

**Figure 2.**
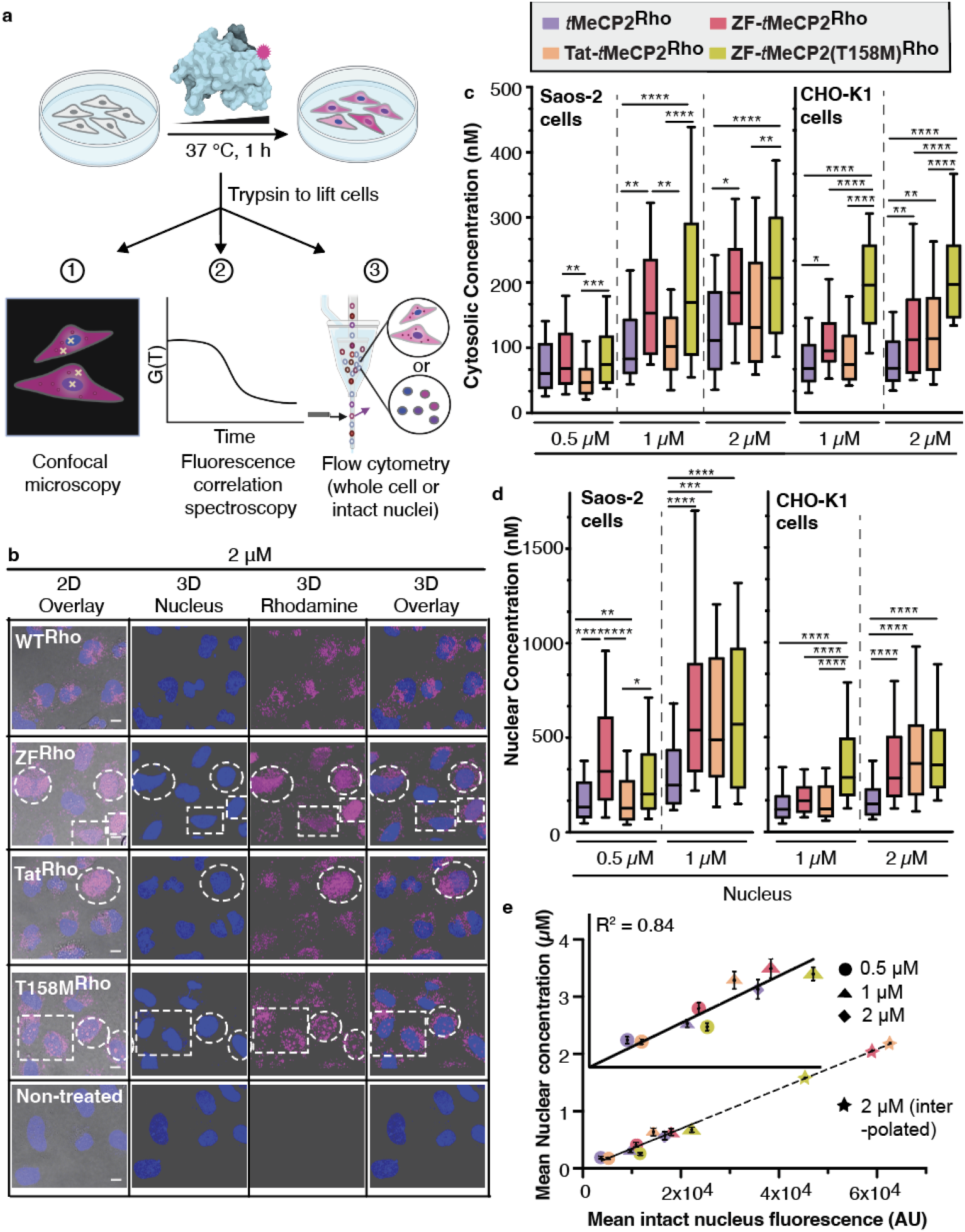
ZF5.3-*t*MeCP2 reaches the nucleus. **a**, Experimental workflow. Saos-2 or CHO-K1 cells were incubated with the indicated *t*MeCP2-Rho variants for 1 hr at 37 °C, 5% CO_2_; Hoechst 33342 was added during the final 5 min of the incubation period to identify the nucleus. After extensive washing and trypsin treatment to remove extracellular and surface-bound protein, cells were analyzed using 2D- and 3D-confocal microscopy and fluorescence correlation spectroscopy (FCS) or flow cytometry. **b**, Representative 3D z-stacking images of Saos-2 cells treated with 2 μM of the indicated *t*MeCP2-Rho variant. Note that all proteins are labeled with Rho to a similar extent (fractional labeling between 20 and 25%). Thus the absence of nuclear fluorescence in cells treated with *t*MeCP2-Rho is not due to differences in labeling efficiency. Cells with nuclear fluorescence are highlighted in the white dash boxes. Scale bar = 10 μm. The results shown are representative of at least two biological replicates. FCS measurements were performed on individual cells by placing the laser focus (represented by the yellow crosshairs in **a**) in either the cytosol, avoiding fluorescent puncta, or the nucleus. The autocorrelation data (**Supplementary Figs. 1-3**) was fitted to a 3D anomalous diffusion equation (cytosol)^42^ or a two-component 3D diffusion equation (nucleus)^44^ using a custom MATLAB script to establish the concentration of each protein in the cytosol (**c**) and nucleus (**d**) of Saos-2 cells or CHO-K1 cells. Center line, median; box limits, 25-75 percentile; whiskers, 10-90 percentile. (n > 30 cells total for each condition from at least two biological replicates, see source data). The intracellular concentrations of the four *t*MeCP2-Rho variants under the same treatment condition and location (e.g. 1 μM in the cytosol) were compared using Brown-Forsythe and Welch ANOVA followed by Dunnett’s T3 multiple comparisons test. ****p ≤ 0.0001, ***p ≤ 0.001,**p ≤ 0.01, *p ≤ 0.05. Each P value is adjusted to account for multiple comparisons (see source data). **d**, Scatter plot depicting the correlation of the mean nuclear fluorescence of intact Saos-2 nuclei measured by flow cytometry (x-axis) and the intra-nuclear concentrations measured by FCS (y-axis). Dots are represented as mean ± SEM. Inset: Data for treatment with 0.5 - 1.0 μM proteins and 2 μM *t*MeCP2-Rho were fitted with a simple linear regression line. The nuclear concentration of ZF-*t*MeCP2-Rho, Tat-*t*MeCP2-Rho and ZF-*t*MeCP2(T158M)Rho in Saos-2 cells treated with 2 μM proteins (stars) were interpolated based on their corresponding nuclear fluorescence values.

We next used fluorescence correlation spectroscopy (FCS) to quantitatively track the cytosolic and nuclear distribution of each *t*MeCP2-Rho conjugate in Saos-2 and CHO-K1 cells (**Fig. 2c-d, Extended Data Table 1, Supplementary Figs. 1-3**). FCS is a single-molecule technique that deconvolutes the time-dependent change in fluorescence in a small cytosolic or nuclear volume to establish both average intracellular concentration as well as diffusion time^42,43^. FCS analysis of Saos-2 cells revealed that all *t*MeCP2-Rho conjugates localize more significantly to the nucleus than the cytosol (**Extended Data Fig. 3a**), as established qualitatively by confocal microscopy (**Fig. 2b, Extended Data Fig. 2, Supplementary Video 1-4)**. Localization to the nucleus is dose-dependent between 500 nM and 1 μM, even for *t*MeCP2-Rho (**Extended Data Fig. 3a**). At low treatment concentrations (0.5 μM), the FCS-determined mean nuclear delivery efficiencies of *t*MeCP2-Rho and Tat-*t*MeCP2-Rho were lower than either ZF-*t*MeCP2-Rho (2.3-fold) or ZF-*t*MeCP2(T158M)-Rho (1.6-fold) (**Fig. 2d, Extended Data Table 1**). At 1 μM, ZF-*t*MeCP2-Rho, ZF-*t*MeCP2(T158M)-Rho, and Tat-*t*MeCP2-Rho reach the nucleus more efficiently (2.0-2.3-fold) than *t*MeCP2-Rho, with localization efficiency increasing in the order: *t*MeCP2-Rho < Tat-*t*MeCP2-Rho < ZF-*t*MeCP2-Rho ∼ ZF-*t*MeCP2(T158M)Rho (**Fig. 2d, Extended Data Table 1**).

**Figure 3.**
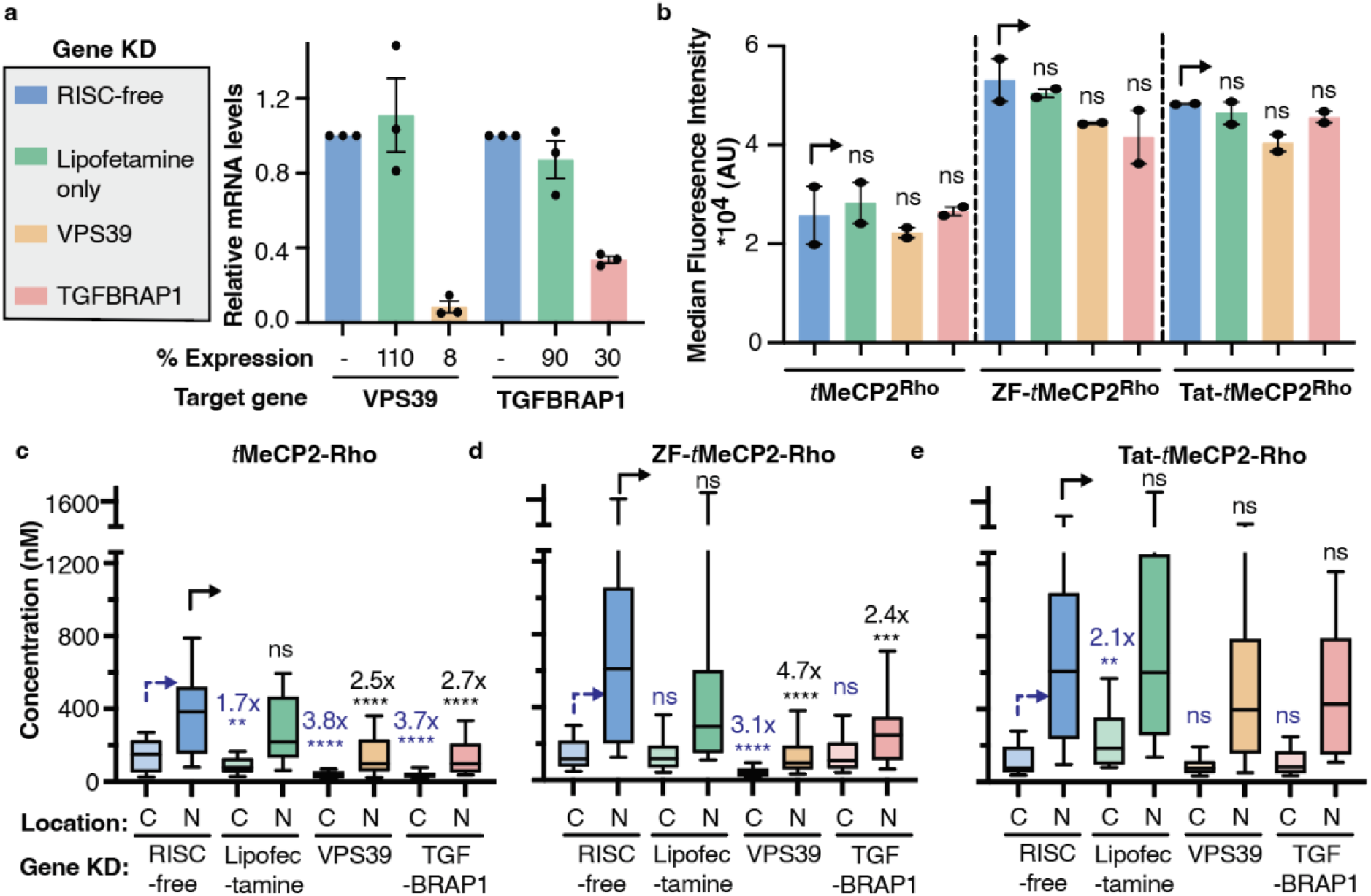
Efficient cellular trafficking of ZF-*t*MeCP2-Rho requires the HOPS complex. **a**, The gene expression level of VPS39 (HOPS-specific subunits) or TGFBRAP1 (CORVET-specific subunits) in each gene knockdown sample was quantified by qPCR. Data are represented as mean ± SEM (n=3). The effect of VPS39 and TGFBRAP1 knockdown on total uptake and cellular access of the indicated *t*MeCP2-Rho variant was analyzed using flow cytometry (**b**) and FCS (**c-e**), respectively. n>4800 each for flow cytometry (mean ± SEM) and n > 20 total for each FCS condition from two biological replicates, see source data. C: cytosol; N: nucleus. Center line, median; box limits, 25-75 percentile; whiskers, 10-90 percentile. For each protein treatment, the median fluorescence intensity values (**b**), cytosolic concentrations (**c-e**) or nuclear concentrations (**c-e**) were compared to that of a non-targeting siRNA (RISC-free) using two tailed unpaired parametric t test with Welch’s correction. ****p ≤ 0.0001, ***p ≤ 0.001,**p ≤ 0.01, *p ≤ 0.05. not significant (ns) for p > 0.05.

We note that while the conjugation of ZF5.3 to *t*MeCP2 resulted in a smaller fold-improvement in nuclear or cytosolic delivery than previously reported examples (improvements between 3^9,11^-32^9^-fold), the mean nuclear concentration of ZF-*t*MeCP2 established in Saos-2 cells after a 1 h incubation with 1 μM protein (709 ± 69 nM) is the highest intracellular concentration yet measured for a delivered protein^9,11^. The high concentration of ZF-*t*MeCP2 that reaches the nucleus may result from the higher intrinsic permeability of *t*MeCP2 itself in comparison to other proteins when evaluated under comparable conditions (argininosuccinate synthetase: 77 ± 30 nM, SNAP-tag: 2 ± 1 nM)^9,11^. While further studies are needed to establish the factors that lead to high intrinsic permeability, we note that *t*MeCP2 is characterized by both a high pI (10.78) and high levels (60%) of intrinsic disorder as judged by CD (**Extended Data Fig. 1c, Supplementary Table 1**).

### Flow cytometry as a high-throughput alternative to FCS for evaluating nuclear delivery *en masse*

The nuclear fluorescence of cells treated with 2 μM ZF-*t*MeCP2-Rho, Tat-*t*MeCP2-Rho and ZF-*t*MeCP2(T158M)-Rho was too high to be reliably measured by FCS^42,43^. Although the total fluorescence of intact cells measured using flow cytometry does not reliably quantify how much material escapes from the endosomal pathway^9^, procedures to isolate and sort nuclei on the basis of fluorescence are well known^45^. We wondered whether the fluorescence of nuclei isolated from cells treated with *t*MeCP2-Rho variants would correlate with the nuclear concentration measured in intact cells using FCS. If such a correlation was observed, then flow cytometry would provide an extremely high-throughput and rapid alternative to FCS for quantifying delivery of fluorescently tagged material to the nucleus.

Thus, we isolated intact nuclei from Saos-2 cells after treatment with 0.5 - 1.0 μM *t*MeCP2-Rho variants and 2 μM *t*MeCP2-Rho and evaluated the nuclear extracts by flow cytometry (**Fig. 2e, Extended Data Fig. 4a-b, Supplementary Figs. 4**). In this concentration range, the mean nuclear fluorescence of intact Saos-2 nuclei measured by flow cytometry correlated linearly (R^2^ = 0.84) with the intra-nuclear concentrations previously measured by FCS for all four Rho-tagged *t*MeCP2 proteins, regardless of overall delivery efficiency (**Fig. 2e)**. A similar linear correlation was observed in CHO-K1 cells (*vide infra*) (**Extended Data Fig. 4c**). This observation suggests that flow cytometry of intact nuclei represents a rapid alternative to FCS for high-throughput analysis of nuclear delivery. It also suggests that at a treatment concentration of 2 μM, the nuclear concentration of ZF-*t*MeCP2-Rho was above 1.5 μM.

**Figure 4.**
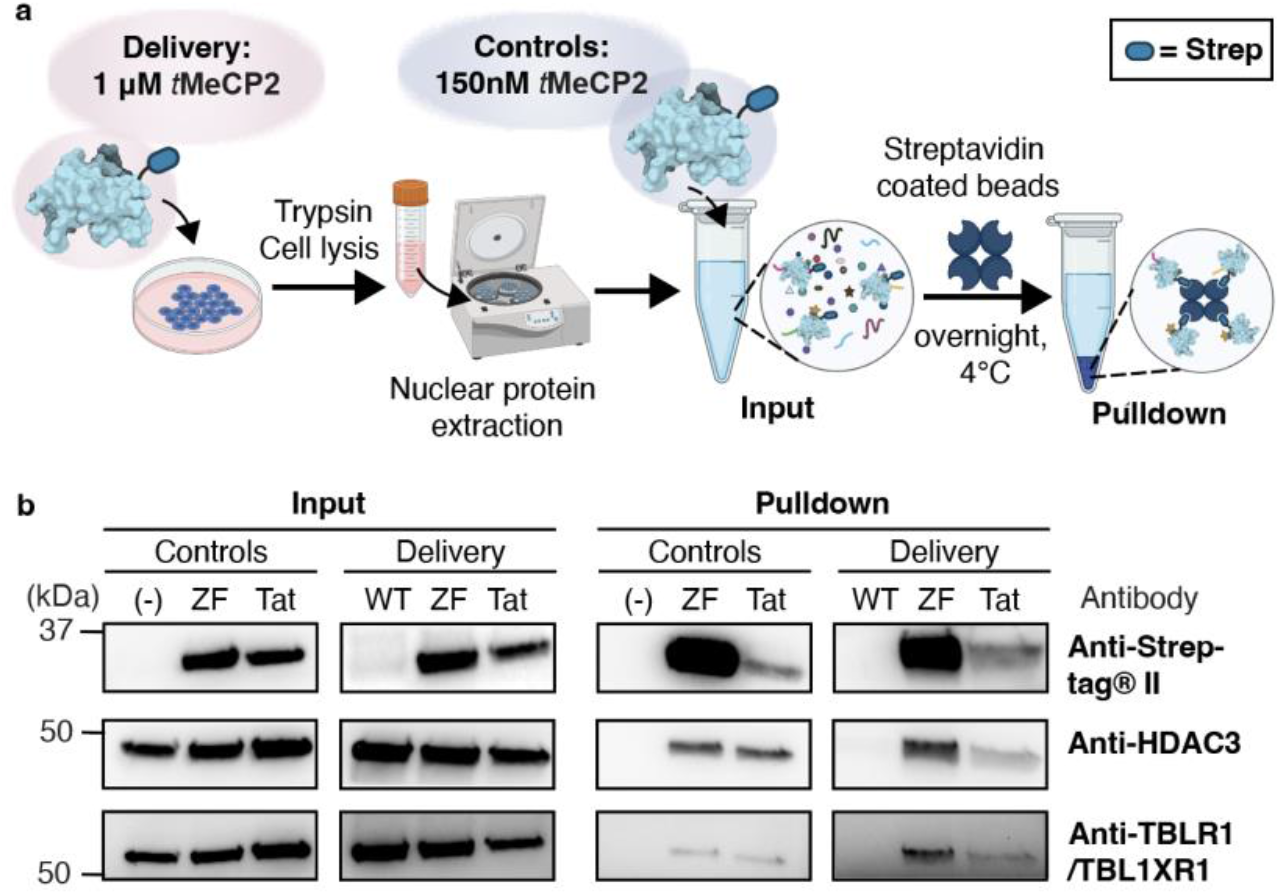
*In cellulo* co-immunoprecipitation assay probing *t*MeCP2 interaction with partner proteins after delivery. **a**, Assay workflow. CHO-K1 cells were treated with 1 μM of each *t*MeCP2 variant at 37 °C with 5% CO_2_ for 1 hr. Three dishes of non-treated cells were incubated under identical conditions as controls. After incubation, cells were lifted, washed, and lysed, and the nuclear proteins were extracted (**Extended Data Fig. 7b**). The nuclear extracts of non-treated cells were doped with 150 nM ZF-*t*MeCP2 or Tat-*t*MeCP2 and then all samples were incubated with streptavidin coated magnetic beads overnight at 4 °C for the pulldown. **b**, Input and pulldown samples were analyzed by Western blot using antibodies against Strep-tagII (IBA 2-1509-001), HDAC3 (CST 85057S) and TBLR1/TBL1XR1 (CST 74499S). The gel results shown are representative of two biological replicates.

To better understand the relationship between overall protein uptake and nuclear delivery, we used flow cytometry to compare whole-cell fluorescence to that of intact isolated nuclei as a function of *t*MeCP2-Rho variant concentration and identity (**Extended Data Fig. 4a, b**). In general, the fluorescence detected in the nuclei of Saos-2 cells was 2.0-5.6-fold lower than the whole-cell fluorescence, but we observed that at 2 μM some cells treated with ZF-*t*MeCP2-Rho and Tat-*t*MeCP2-Rho had nuclear fluorescent intensities as high as those of the whole cell, suggesting the majority of the protein within these cells had trafficked to the nucleus (blue dashed box in **Extended Data Fig. 4a**). Comparing the mean nuclear fluorescence to the mean whole-cell fluorescence (**Extended Data Fig. 4b**), there is a general trend of dose-dependent increase in the fraction that travels to the nucleus. At treatment concentrations below 2 μM, the fraction of *t*MeCP2-Rho variants that reached the nucleus was low (17-30%), regardless of both concentration and whether or not *t*MeCP2 was fused to ZF5.3, Tat, ZF5.3(T158M). At 2 μM, however, the fraction of *t*MeCP2-Rho variant that reached the nucleus was significant for ZF-*t*MeCP2-Rho (52%), Tat-*t*MeCP2-Rho (51%), as well as ZF-*t*MeCP2(T158M)-Rho (43%). In preparation for function studies (*vide infra*), we also evaluated the trafficking of each *t*MeCP2-Rho variant to the cytosol and nucleus of CHO-K1 cells using confocal microscopy (**Extended Data Fig. 5**), flow cytometry (**Extended Data Fig. 4c, d**), and FCS (**Fig. 2c-d, Extended Data Table 1, Extended Data Fig. 3b,c, Supplementary Fig. 3**). These results largely mirrored the results obtained in Saos-2 cells, although overall the protein concentration in the nucleus was 1.8-3.8 fold lower than observed in Saos-2 cells. Similar cell type-to-cell type variations in delivery efficiency have been observed before among HeLa cells, SK-HEP1 cells, and Saos-2 cells^9,11^. Nevertheless, these results provide confidence that ZF5.3 and Tat improve by 2.0-2.5 fold the nuclear delivery of *t*MeCP2-Rho in two model cell lines.

**Fig. 5.**
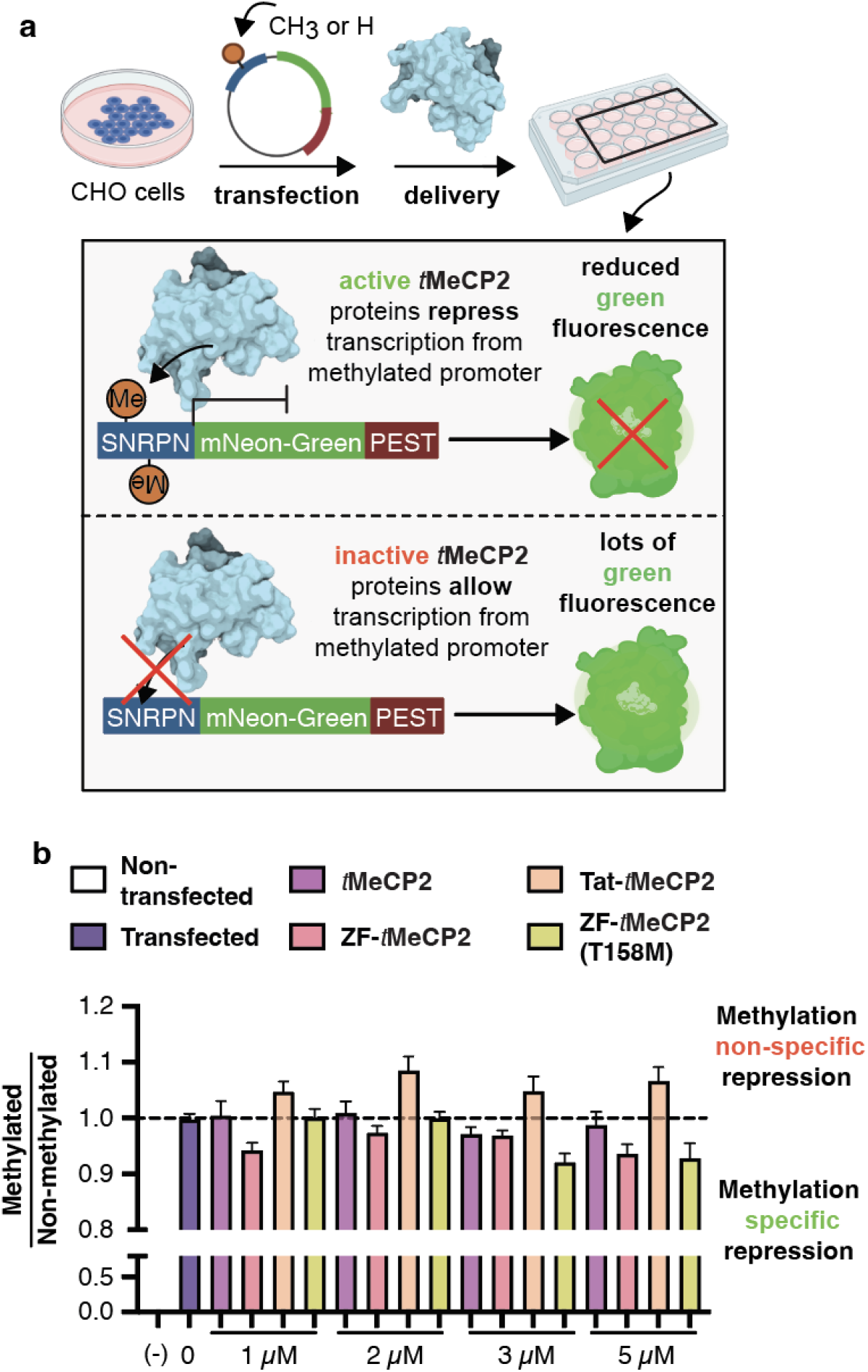
**Transcription repression assay probing** *t***MeCP2 functionality *in cellulo*.** **a**, Scheme illustrating assay design. CHO-K1 cells were transiently transfected with a plasmid (methylated or non-methylated) encoding mNeonGreen fluorescent protein under the control of the small nuclear ribonucleoprotein polypeptide N (SNRPN) promoter. Cells were first incubated with 1 μM *t*MeCP2 variants for 1 h at 37 °C, 5% CO_2_, and exchanged into growth media for 1 h at 37 °C, 5% CO_2_ waiting for the change in transcription to occur before they were lifted and analyzed using flow cytometry. Cells that were successfully transfected and exhibited green fluorescence higher than the background were selected to obtain the mean fluorescence intensity (MFI) under different protein treatments. The assay was performed in triplicates and more than three biological replicates were studied. **b**, The selectivity of four *t*MeCP2 variants was evaluated by dividing the average normalized MFI after delivery in CHO-K1 cells transfected with methylated plasmids by that with non-methylated plasmids. Scores are normalized against non-treated transfected cells. Data are represented as mean ± SEM. Each sample comprised 130 μL (at least 50,000 cells) and at least three technical and biological replicates at each condition, see source data.

### ZF-*t*MeCP2, but not Tat-*t*MeCP2, is intact when it reaches the nucleus

Although fluorescence-based methods offer a good initial assessment of protein delivery efficiency, they must always be accompanied by biochemical studies to ensure that the fluorescent material being followed is intact^9,11,42^. To establish the extent to which the measured FCS values represent the concentrations of intact *t*MeCP2 proteins, we devised a workflow (**Extended Data Fig. 6a**) to stringently separate and isolate the cytosolic and nuclear fractions of Saos-2 cells after 1 hr treatment with 1 μM *t*MeCP2, ZF-*t*MeCP2, or Tat-*t*MeCP2 at 37 °C. These extracts were analyzed using Western blots and an anti-Strep-tag antibody. Non-treated cells were subject to the same workflow and doped with 150 nM purified *t*MeCP2, ZF-*t*MeCP2, or Tat-*t*MeCP2 to calibrate the Western blots (**Extended Data Fig. 6b**). Bands corresponding to both intact ZF-*t*MeCP2 and Tat-*t*MeCP2 are evident in the nuclear supernatant. We note that although the nuclear delivery efficiencies of ZF-*t*MeCP2-Rho and Tat-*t*MeCP2-Rho determined by FCS were comparable (**Fig. 2d**), Western blot analysis suggests that the concentration of intact ZF-*t*MeCP2 in the nucleus exceeds that of Tat-*t*MeCP2 by a significant margin. No band corresponding to *t*MeCP2 itself was observed in the nuclear supernatant, indicating that this protein was either degraded or did not enter cells at a concentration high enough to be detected. Western blot analysis of cellular extracts with antibodies recognizing highly expressed endocytic (EEA1, LAMP1, Rab7) and cytosolic (tubulin) proteins confirmed that nuclear fractions were free of detectable cytosol and endosome contaminations (**Extended Data Fig. 6c**).

To further characterize the integrity of ZF-*t*MeCP2 delivered to the nucleus, we enriched the nuclear supernatant for strep-tagged proteins, treated the enriched sample with trypsin, and subjected the digest to LC-MS/MS analysis (**Extended Data Fig. 6d, Supplementary Fig. 5**). More than 65% of the ZF-*t*MeCP2 sequence was detected, including fragments at the N- and C-terminus. The observation of N-terminal fragments after enrichment with a C-terminal strep tag provides further evidence that the ZF-*t*MeCP2 delivered to the nucleus is predominantly intact.

### ZF-*t*MeCP2 accesses a HOPS-dependent portal for endosomal release

Two multisubunit tethering complexes play important roles in the endosome maturation pathway. A class C core vacuole/endosome tethering (CORVET) complex facilitates the fusion of Rab5 positive early endosomes while the fusion of Rab7 positive late endosomes to lysosomes requires the homotypic fusion and protein sorting (HOPS)-tethering complex^46,47^. Previous mechanistic studies indicate that efficient cytosolic and nuclear trafficking of ZF5.3 relies on the HOPS complex, but not the analogous CORVET complex^10^. We thus sought to investigate if this dependence also held for the ZF5.3 conjugate of *t*MeCP2.

We used siRNAs to knock down an essential and unique subunit of either HOPS (VPS39) or CORVET (TGFBRAP1) in Saos2 cells, using a non-targeting siRNA (RISC-free) and lipofectamine only treatment as controls (**Fig. 3a**). The cells were then treated with 1 μM *t*MeCP2-Rho, ZF-*t*MeCP2-Rho, or Tat-*t*MeCP2-Rho for 1 h and analyzed by flow cytometry (**Fig. 3b**) and FCS (**Fig. 3c-e)**. Total cellular uptake was affected minimally if at all by any gene knockdown (**Fig. 3b**). Knockdown of VPS39 led to a significant decrease in the concentration of ZF-*t*MeCP2-Rho that reached the cytosol (3.1-fold) or nucleus (4.7-fold) (**Fig. 3d**); these fold changes are consistent with those previously observed for ZF5.3 alone^10^. Interestingly, knockdown of TGFBRAP1 also resulted in a significant (albeit smaller) decrease in the concentration of ZF-*t*MeCP2-Rho that reached the nucleus (2.4-fold). Even the trafficking of *t*MeCP2 itself was affected by the knockdown of both VPS39 and TGFBRAP1 (**Fig. 3c**). It is well known that depletion of TGFBRAP1 and VPS39 can affect trafficking in a cargo-dependent manner^48^. The fact that delivery of *t*MeCP2 itself is CORVET and HOPS dependent provides one explanation for the high intrinsic permeability of this nuclear protein and deserves further study. Thus the improved nuclear delivery of ZF-*t*MeCP2 may result because it accesses both HOPS-dependent and CORVET-dependent portals. Notably, the only *t*MeCP2 conjugate whose delivery to the cytosol and nucleus was unaffected by knockdown of either VPS39 or TGFBRAP1 was Tat-*t*MeCP2 (**Fig. 3e**). This result suggests that Tat-*t*MeCP2 gains entry into cells, albeit less efficiently (**Extended Data Fig. 6b**), *via* a different subpopulation of endosomes^48^ or one or more non-endosomal pathways^49^.

### Delivered ZF-*t*MeCP2 interacts with partner proteins in the NCoR/SMRT complex

Next we explored the function of *t*MeCP2 proteins delivered to the nucleus of CHO-K1 cells, which express low levels of endogenous MeCP2 (**Extended Data Fig. 7a**). If functional *t*MeCP2 proteins reach the nucleus, then they should interact with and sequester the essential subunits of the core NCoR/SMRT complex^6,34,36^ (NCoR1, HDAC3, and TBL1/TBLR1) upon immunoprecipitation, as observed *in vitro* in lysates (**Fig. 1c**). To test this hypothesis, CHO-K1 cells were treated with 1 μM *t*MeCP2, ZF-*t*MeCP2, or Tat-*t*MeCP2 for 1 hr at 37 °C. Nuclear proteins were rigorously isolated (**Extended Data Fig. 7b)** and incubated with streptavidin-coated magnetic beads to sequester strep-tagged *t*MeCP2 proteins and the proteins with which they interact (**Fig. 4a**). Non-treated cells were subject to the same workflow and doped with 150 nM purified *t*MeCP2, ZF-*t*MeCP2, and Tat-*t*MeCP2. Western blot analysis confirmed that the input nuclear fractions used for immunoprecipitation were free of detectable cytosolic (tubulin) and endosomal (EEA1, Rab7, LAMP1) contaminants (**Extended Data Fig. 7c**) and contained a higher amount of intact ZF-*t*MeCP2 than Tat-*t*MeCP2, in accordance with data in Saos-2 cells in **Extended Data Fig. 6**. Intact *t*MeCP2 could not be detected by Western blot of the nuclear extracts. Equal amounts of HDAC3 and TBLR1/TBL1XR1 were also detected in all input nuclear lysates.

Examination of the Western blots after immuno-precipitation show that both ZF-*t*MeCP2 and Tat-*t*MeCP2 sequester HDAC3 and TBLR1 from nuclear lysates in accord with their effective concentration; more ZF-*t*MeCP2 reaches the nucleus intact and as a result more HDAC3 and TBLR1 are sequestered (**Fig. 4b**). We note that the large decrease in the intensity of the Tat-*t*MeCP2 bands from input to pulldown emphasizes its sensitivity to degradation during the overnight incubation. The NcoR level was too low to be detected in CHO-K1 cells (**Extended Data Fig. 7a**). We conclude that ZF-*t*MeCP2 enters the cell nucleus and interacts more productively with partner proteins than either *t*MeCP2 or Tat-*t*MeCP2.

### Delivered ZF-tMeCP2 selectively represses transcription from methylated DNA

In the nucleus, MeCP2 acts as a bridge to deliver the NCoR/SMRT complex to methylated promoters; this recruitment represses transcription^23^. We devised a flow cytometry assay to evaluate whether delivered *t*MeCP2 variants that reach the nucleus preferentially repress transcription of methylated over non-methylated reporter genes (**Fig. 5a**). Briefly, CHO-K1 cells were transfected with a methylated or non-methylated plasmid encoding mNeonGreen fluorescent protein under the control of the small nuclear ribonucleoprotein polypeptide N (SNRPN) promoter. MeCP2 binds to the methylated form of SNRPN to downregulate downstream genes^50,51^. A short signal sequence (PEST) was encoded at the C-terminus of mNeonGreen to promote its turnover and improve assay sensitivity^52^. Functional *t*MeCP2 variants that reach the nucleus should selectively repress transcription from cells transfected with the methylated SNRPN promoter, leading to less mNeonGreen fluorescence relative to controls. By contrast, delivery of a trace, non-functional, or nonspecific *t*MeCP2 variant will either not repress transcription or do so without selectivity for the methylated promoter.

To evaluate mNeonGreen expression, CHO-K1 cells transfected with methylated or non-methylated plasmid DNA were treated for 1 hr at 37 °C with 1-5 μM *t*MeCP2, ZF-*t*MeCP2, Tat-*t*MeCP2 or ZF-*t*MeCP2(T158M) and the fluorescence emission at 530 ± 30 nm was monitored using flow cytometry (**Supplementary Fig. 4**). Cells treated with *t*MeCP2 itself displayed high mNeonGreen fluorescence levels regardless of promoter methylation state (**Extended Data Fig. 8**). By contrast, cells treated with ZF-*t*MeCP2, Tat-*t*MeCP2, as well as ZF-*t*MeCP2(T158M) all showed dose-dependent decreases in mNeonGreen fluorescence. The highest levels of methylation-specific transcriptional repression was observed in cells treated with 5 μM ZF-*t*MeCP2, Tat-*t*MeCP2, as well as ZF-*t*MeCP2(T158M), although measurable effects were seen at concentrations as low as 2 μM. In general, the levels of transcriptional repression were lower in cells treated with Tat-*t*MeCP2.

These data are in line with the protein nuclear concentrations (**Fig. 2d**) and diffusion time and DNA binding kinetics measured by FCS (**Extended Data Table 2**). FCS is useful not only for measuring the concentration of a fluorescently tagged protein or macromolecule within the cytosol or nucleus, but also for determining its intracellular diffusion time (**τ**_diff_)^42,43^. Fitting the autocorrelation curves obtained from intra-nuclear measurements in CHO-K1 cells with a two-component 3D diffusion equation^53^ identified a population of *t*MeCP2-Rho variants that diffuses freely in the nucleoplasm (fast fraction, *F*_fast_) and a second population that diffuses slowly (slow fraction, *F*_slow_), presumably because it is bound to DNA (**Supplementary Fig. 3, Extended Data Table 2**). At 1 μM, the fraction of ZF-*t*MeCP2 diffuses slowly (*F*_slow_ = 26.3%) is higher than that of ZF-*t*MeCP2(T158M) (16.3%) in accord with relative methylated DNA affinities determined *in vitro* (**Fig. 1b**) and in cells^33^. Thus the higher nuclear concentration of ZF-*t*MeCP2(T158M) is counterbalanced by the low DNA binding population; the result is no significant transcriptional repression (**Extended Data Fig. 8)**. At 2 μM, ZF-*t*MeCP2, Tat-*t*MeCP2, as well as ZF-*t*MeCP2(T158M) all reached the nucleus at significantly higher levels than *t*MeCP2 (**Fig. 2d**) and show higher levels of DNA binding, with values of *F*_slow_ of 24.9%, 33.7% and 25.6%, respectively (**Extended Data Table 2**); the result is observable transcriptional repression.

Differences in transcriptional repression are most apparent when the ratio of mNeonGreen expression in cells transfected with methylated *vs*. non-methylated promoters are compared (**Fig. 5b**). As expected, no selective repression is observed in cells treated with *t*MeCP2. Tat-*t*MeCP2 treatment led to higher levels of mNeonGreen repression in cells transfected with a non-methylated promoter, as suggested by the *in vitro* DNA binding results (**Fig. 1b**); Tat alone possesses high non-specific DNA binding affinity (K_d_ = 126 nM)^54^. The highest levels of methylation-dependent transcriptional repression were observed in cells treated with ZF-*t*MeCP2 and ZF-*t*MeCP2(T158M). As a previous study suggested, MeCP2(T158M) is capable of binding to methylated DNA in a protein level-dependent manner^55^. At low treatment concentrations (1 - 2 μM), most ZF-tMeCP2(T158M) was sequestered in the nucleolus, so the effective nuclear concentration of ZF-tMeCP2(T158M) is not high enough to rescue its reduced methylated DNA binding affinity. As more protein reaches the nucleus at 5 μM treatment (∼2 μM interpolated based on its corresponding nuclear fluorescence value in **Extended Data Fig. 3c**), a higher level of the methylated promoter is bound and repression is observed. Taken together, these data indicate that the fusion of ZF5.3 to tMeCP2 does not interfere with the protein’s selectivity towards methylated DNA. ZF-*t*MeCP2 reaches the nucleus at a concentration high enough to observe methylation-dependent transcription repression. Notably, although fluorescent detection implies relatively comparable nuclear delivery of ZF-*t*MeCP2 and Tat-*t*MeCP2, the latter is largely degraded, impairing its ability to regulate downstream transcriptions.

## Discussion

Efficient protein delivery remains a significant and unmet challenge in an era exploding with novel protein-based therapeutic strategies. Almost all FDA-approved biologics are delivered *via* injection, and operate within serum, on cell surfaces, or within endosomal vesicles. The impact of protein-based strategies would expand exponentially if proteins could reliably evade endosomal degradation, escape endosomal vesicles, and remain functional. Yet progress towards this goal has been exceptionally slow. Most delivery strategies are inefficient^9^, rely on degradation-prone molecular scaffolds^26,29^, operate *via* undefined or multifarious mechanisms^56^, and detect activity using amplified or indirect assays^9,57^.

In this work, we address these challenges by using the stable mini-protein ZF5.3 to hijack the endosomal maturation machinery and guide the methyl-CpG-binding-protein 2 (MeCP2) into the nucleus at a therapeutically relevant concentration. Rigorous analyses verify the absence of significant nuclear degradation and the presence of MeCP2-specific activity. Multiple independent approaches, including intact nuclear flow (INF), a novel and general application of flow cytometry, quantitatively assess delivery in a non-amplified and high-throughput manner. Finally, the remarkable level of nuclear delivery by ZF-*t*MeCP2 enables its potential application to reverse Rett syndrome phenotypes; a modest dose to Saos-2 cells (1 h, 1 μM) results in a nuclear concentration of 2.1 × 10^5^ molecules/cell, which is within the range of endogenous MeCP2 in HEK293^24^ and NeuN positive neurons from mature mouse brain^1^ (1.6 × 10^5^ and 160 × 10^5^ molecules/cell, respectively). This delivered concentration is two orders of magnitude higher than a previously reported Tat-MeCP2 conjugate that could rescue some RTT-related symptoms^26,27,29^.

Progress in direct protein delivery has also been slowed by an insufficient understanding of how non-native cargo traffics across endosomal membranes and into the cytosol. Although endosomal escape mechanisms used by certain viruses, such as influenza A^58^, have been studied in detail, those exploited by non-viral agents – even lipid nanoparticles (LNP) used to deliver FDA-approved mRNA vaccines – remain incompletely characterized. Most improvements in delivery are empirical and modest, and the involvement of endogenous protein factors has not been rigorously excluded. The highly efficient trafficking of ZF5.3 itself into the cytosol demands an intact and functional homotypic fusion and protein-sorting (HOPS) complex^10^, a membrane-tethering complex that fuses Rab7+ endosomes during the natural process of endosomal maturation. HOPS may engage directly with ZF5.3 to promote escape during vesicle fusion; it may also promote trafficking into intraluminal vesicles as a prerequisite to endosomal escape. The observation that the high nuclear delivery efficiency of ZF5.3-*t*MeCP2 depends on both HOPS and CORVET, a related endosomal maturation machine that operates earlier along the endocytic pathway, favors the former explanation and deserves further study, especially as HOPS and CORVET share 4 of 6 subunits in common^46^. Identification of those molecular features that promote productive interaction with HOPS and/or CORVET could improve the efficiency of other delivery strategies, including lipid nanoparticles, whose efficiency remains < 10%^59^.

## Methods and Materials

### Plasmid construction

We initially attempted to express and purify full-length MeCP2 as a C-terminal fusion with *ZF5*.*3 in Escherichia coli*, but the yield was low and could not be optimized. The DNA sequence for *t*MeCP2(ΔNC) was obtained by deleting the N-terminal (residues 13-71) and the C-terminal (residues 313-484) sequences of mouse tMeCP2_e2 (protein sequence identifier: Q9Z2D6-1). The codon-optimized gblock encoding ZF5.3-*t*MeCP2(ΔNC)-LPETGG-Strep was purchased from Integrated DNA Technologies (IDT). A GSG linker was added between ZF5.3 and tMeCP2(ΔNC) and a GSSGSSG linker was inserted before the C-terminal LPETGG-Strep tag. A pET His6 MBP TEV LIC cloning vector (Addgene Plasmid #29656) was digested using restriction enzymes XbaI and BamHI to remove the His6-MBP-TEV portion and the synthetic gblocks containing complementary overhangs were incorporated into the pET vector using Gibson assembly. To generate *t*MeCP2(ΔNC)-LPETGG-Strep or Tat-*t*MeCP2(ΔNC)-LPETGG-Strep, the ZF5.3-*t*MeCP2(ΔNC)-LPETGG-Strep plasmid was double digested with XbaI and SacII respectively to remove the ZF5.3 segment, and ligated back using Gibson assembly with double-stranded fragments containing corresponding sequences. Point mutations (T158M, P302L) in the protein were generated by Q5 site-directed mutagenesis. The identity of all plasmids were confirmed by Sanger and whole plasmid sequencing. Relevant DNA and protein sequences are listed in **Supplementary Table 3**

### Protein expression and purification

All *t*MeCP2 variants (with and without labeled fluorophore) are purified as described in **Supplementary Methods 1**.

### Circular Dichroism

Circular Dichroism measurements were performed following a previously published protocol.^60^ Wavelength-dependent circular dichroism spectra were recorded using an AVIV Biomedical, Inc. (Lakewood, NJ) Circular Dichroism Spectrometer Model 410 at 25 °C in a 0.1 cm pathlength quartz cuvette. CD spectra were collected at ∼12 μM protein concentration in 20 mM HEPES, 300 mM NaCl, 10 % glycerol pH 7.6 between 300 and 200 nm at 1 nm intervals with an averaging time of 5 seconds. A separate wavelength spectrum for storage buffer alone was performed to verify no interfering buffer signal. Raw ellipticity values were converted to mean residue ellipticity using the equation

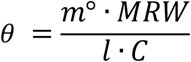

where *θ* = mean residue ellipticity in deg▪cm^2^ ▪dmol^−1^, *m*° = raw ellipticity values in millidegrees, *MRW* = mean residue weight (calculated as the molecular weight divided by the number of backbone amides), *l* = path length of the cuvette in millimeters, and *C* = concentration of protein in mg mL^−1^. The protein secondary structure was predicted using the webserver BeStSel (http://bestsel.elte.hu/index.php).

### Fluorescence polarization

Samples for fluorescence polarization (FP) studies were prepared by mixing assay buffer (25 mM Tris-HCl, 6% glycerol, 100 μg/mL BSA, 0.1 mM EDTA, and 0.1 mM TCEP, 250 mM KCl, pH 7.6), different *t*MeCP2 variants with final concentrations ranging from 1 nM to 5 μM and 20 nM DNA probe. The DNA probes carrying an N terminal 6-fluorescein^61^ (methylated or non-methylated, **Supplementary Table 3**) were synthesized by annealing equimolar of complementary single strands derived from a 22 bp DNA segment of mouse BDNF promoter IV (purchased from IDT) at 95 °C for 2 min, followed by 60 °C for 10 min and cooled to 4 °C^62^. The samples were incubated for 30 min at room temperature to reach equilibrium and added to a 96 well half area solid black plate (Corning #3993, 50 μL per well).

FP measurements were performed using a SpectraMax M5 plate reader with an excitation wavelength at 480 nm and an emission window of 515 to 520 nm. Measurements from three wells were averaged for each determination. To calculate K_D_, the data were fitted by least squares regression to an equilibrium binding equation based on the Langmuir model with modifications explicitly considering the total DNA concentration^39^ using GraphPad Prism 9.

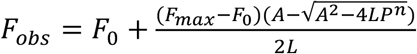

where 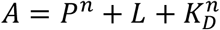. In this equation *F*_0_ is the background FP value of the DNA probe when no protein was added; *F*_*obs*_is the FP detected by the plate reader for each sample; *F*_*max*_ is the maximum FP at saturation; *P* and *L* represent the total protein and DNA concentration, respectively; *K*_*D*_ is the equilibrium dissociation constant; and *n* is the Hill coefficient. While previous studies reported both MBD^39^ and full-length tMeCP2^38,63^ can bind cooperatively to DNA, the high salt concentration used in the assay buffer suppressed binding cooperativity and preserved the specificity for methylated DNA^63^. The Hill coefficients fitted for all conditions in this experiment were close to 1.

### Cell culture

Saos-2 cell stock was purchased from the American Type Culture Collection (ATCC). NIH3T3, HeLa and CHO-K1 cell stocks were purchased from UC Berkeley Cell Culture Facility. Saos-2 cells were cultured in McCoy’s 5A medium (Hyclone) with phenol red containing 15% fetal bovine serum (FBS), penicillin and streptomycin (P/S, 100 units/mL and 100 μg/mL respectively), 1mM sodium pyruvate (Gibco), 2 mM GlutaMax (Gibco). NIH3T3 and HeLa cells were cultured in Dulbecco’s Modified Eagle’s Medium (DMEM, Gibco) with 10% FBS and P/S. CHO-K1 cells were cultured in F12 Nutrient Mixture (Ham) media (Gibco) with L-Glutamine, 10 % FBS and P/S. All cell cultures were incubated at 37 °C, in 5% CO_2_.

### *In vitro* pull-down assay

Nuclear lysate of NIH3T3 cells were obtained as described in **Supplementary Methods 2**. 200 μL of the nuclear lysate was mixed with 1.5 μM of purified *t*MeCP2 variants overnight at 4°C in the coIP buffer^36^ (20 mM HEPES, pH 7.6, 10 mM KCl, 150 mM NaCl, 1 mM MgCl_2_, 0.1% Triton X-100 (vol/vol), protease inhibitors (Roche), 15 mM BME). On the next day, to pull down *t*MeCP2 protein complexes, MagStrep “type3” XT magnetic beads (IBA 2-4090-002) following the manufacturer’s note were used. The beads were pre-equilibrated with coIP buffer and incubated with the mixture of nuclear lysate and *t*MeCP2 for 45 min at RT on a rotating wheel. After separating the beads from the unbound supernatant using a magnetic separator, the beads were washed with 200 μL coIP buffer two times (quickly vortex and centrifuge). The pull-down proteins were eluted with 10 μL 5x SDS-PAGE loading dye and 10 μL milliQ water at 95 °C for 3 min. The input and pulldown samples were analyzed by western blot using primary antibodies against Strep-tagII (IBA 2-1509-001), HDAC3 (CST 85057S), TBLR1/TBL1XR1 (Abcam ab190796), NCoR1 (CST 5948S), and secondary antibody HRP-linked Anti-rabbit IgG (CST 7074S).

### Fluorescence-based assay for protein delivery

The fluorescence-based experiments (confocal microscopy, fluorescence correlation spectroscopy and whole-cell flow cytometry) to evaluate protein delivery were performed according to previously published protocols^42^. For delivery into Saos-2 cells, 500 μL *t*MeCP2 variants diluted in McCoy’s 5A medium (–FBS, –phenol red) to final concentrations (0.5 - 2 μM) were added to the cells plated in a 6-well plate to incubate for 1h at 37°C, 5% CO_2_. 300 nM of the nuclear stain Hoechst 33342 was added to cells 5 min prior to the end of incubation time.

For experiments using CHO-K1 cells, a similar protocol was followed except: the cells were plated in DMEM media supplemented with 1x non-canonical amino acid (+10% FBS, -phenol red) on day 1. On day 2, 1-5 μM *t*MeCP2 variants were diluted in DMEM media supplemented with 1x non-canonical amino acid (-phenol red) for delivery. DMEM media supplemented with 1x non-canonical amino acid (+10% FBS, -phenol red) was used to quench the trypsin reaction and rinse the 6-well plate. Cells were resuspended in DMEM media supplemented with 1x non-canonical amino acid (–phenol red) for confocal microscopy and fluorescence correlation spectroscopy studies.

### Confocal Microscopy and Fluorescence Correlation Spectroscopy (FCS)

Confocal microscopy and FCS experiments were performed using a STELLARIS 8 confocal microscope (Leica) with a Hybrid HyD X detector, and an HC PL APO 63x/1,20 W motCORR CS2 water (63W) immersion objective (Leica). The correction collar of the objective was adjusted based on the thickness of an 8-well chambered coverglass prior to experiments. The fluorophore was excited at 561 nm and the fluorescence filter was set to 570-660nm. The pinhole of the 561 nm laser was set to 1 AU. At the beginning of the experiments, the imaging chamber was equilibrated to 37°C, 5% CO_2_. Before each FCS measurement, a confocal image of the cells was obtained and the laser was directed to either the cytosol or nucleus of the cells to obtain FCS data in discrete cellular locations. Areas around the punctate fluorescence of endosomes were avoided. All FCS measurements were collected in ten consecutive five-second time intervals. A minimum of 20 cells was measured per replicate, and data from at least two biological replicates were collected for each condition. To measure the diffusion coefficients of *t*MeCP2-Rho variants *in vitro*, proteins were diluted in DMEM (25 mM HEPES, -phenol red) to 100 nM fluorophore concentration, and their autocorrelation data were collected at 37°C (10 repeats, 5s intervals). Because some phototoxicity was observed during FCS in CHO-K1 cells, besides keeping the cells at 37°C, 5% CO_2,_ we avoided shining light on a single cell two times by measuring the cytosol, and nuclear concentrations in separate cells.

The cytosolic and nuclear concentrations of fluorescently tagged *t*MeCP2-Rho variants in cells were calculated by fitting the measured data to a 3D anomalous diffusion equation (cytosol) and a two-component 3D diffusion equation (nucleus). We further adjusted the calculated concentrations based on the fluorophore labeling efficiency of each protein to obtain the true intracellular concentrations (**Supplementary Note 1**).

### Flow cytometry

Flow cytometry measurements were performed at room temperature with an Attune NxT flow cytometer. The fluorophore Rho was excited with a laser at 561 nm, and the emission filter was set at 585 ± 16 nm. For whole-cell flow cytometry, at least 10,000 cells were analyzed for each sample and at least three biological replicates were measured for each condition. To isolate the intact nuclei, cells treated with *t*MeCP2 variants were washed and trypsinized as described above. After centrifugation at 200 g for 3 min, the pellet was resuspended in 1 mL DPBS and 200 μL was saved for whole-cell flow cytometry analysis. The rest of 800 μL was pelleted at 200 g for 3min, resuspended with precooled 1 mL isotonic sucrose buffer (290 mM sucrose, 10 mM imidazole, pH 7.0, 1 mM DTT, and 1 cOmplete protease inhibitor cocktail (Roche) per 10 mL buffer) and transferred to a 1.5 mL microcentrifuge tube. After centrifuging at 10,000 g, 4 °C for 1 min, the pellet was resuspended in 200 μL of isotonic sucrose buffer + 0.1% NP-40 and centrifuged at 1000 g, 4 °C for 10 min. The resulting intact nuclei pellet was resuspended in 200 μL PBS. 20 μL sample was mixed with Trypan blue to check the intactness of the isolated nuclei under an inverted microscope while the rest was analyzed by flow cytometry. The collected data were analyzed using FlowJo software (version 10.8.1, FlowJo, LLC).

### RNAi

siRNA-mediated knockdown was performed as described previously^10^ using Lipofectamine RNAiMAX transfection reagent (Invitrogen) and siRNA (100 nM, Dharmacon). All siRNA used was listed in **Supplementary Table 3**. Saos-2 cells were transfected for 4 hours and grew in McCoy’s 5A medium (2 mL/well, 15% FBS, 1 mM sodium pyruvate, 2 mM GlutaMax, -phenol red, -P/S) for 72 h at 37 °C, 5% CO2 for optimal gene knockdown. The effect of gene knockdown was studied using flow cytometry and FCS as described above.

### RT-qPCR

The degree of siRNA-mediated gene knockdown after 72 h was evaluated using RT-qPCR as described previously^10^. For RT-qPCR, cDNA was amplified using 150 nM gene-specific primers (IDT, PrimeTime qPCR primers) with SsoFast EvaGreen Supermix (Bio-Rad) on a Bio-Rad CFX96 real-time PCR detection system. Three biological replicates were evaluated. Each run includes the siRNA-targeted gene, GAPDH as the reference gene and their respective negative controls (no reverse transcriptase and no template). Each sample was run in triplicate. Primers used for reverse transcription into cDNA and qPCR were listed in **Supplementary Table 3**.

### Cytosolic fractionation to determine protein intactness

The cellular fractionation experiment was adapted from previous reports^9–11^ with modifications to ensure pure nuclei isolation. Saos-2 cells were grown to ∼2.5 × 10^6^ cells in a 100 mm dish in McCoy’s 5A medium (+15% FBS, -phenol red). For protein delivery, *t*MeCP2-Strep variants diluted in clear McCoy’s media without FBS to final concentration of 1 μM were incubated with cells at 37 °C with 5% CO_2_ for 1 h. One dish of cells with the same clear McCoy’s media added was included as a non-treated control. After incubation, cells were lifted, washed, and lysed as previously described.^11^ The homogenized cell lysate was centrifuged for 10 min at 800 g, 4 °C. The crude supernatant was re-centrifuged at 1200 g, 4 °C for 10 min. The resulting supernatant was transferred to a polycarbonate ultracentrifuge tube and centrifuged at 350 kg for 30 min (TL-100; Beckman Coulter, TLA-100 rotor (20 × 0.2 mL) to isolate the cytosolic fraction. The crude nuclei pellet was washed in 500 μL of isotonic sucrose buffer (290 mM sucrose, 10 mM imidazole, pH 7.0, 1 mM DTT, and 1 cOmplete protease inhibitor cocktail (Roche)) supplemented with 0.15% NP-40. After incubating on ice for 10 min, the mixture was centrifuged at 1200 g, 4 °C for 10 min. The resulting pure nuclear pellet was first resuspended with 20 μL milliQ water and 2 μL benzonase (Sigma, 71206) at room temperature for 20 min. 80 μL high salt extraction buffer^64^ (20 mM HEPES, pH 7.6, 1.5 mM MgCl_2_, 420 mM NaCl, 0.2 mM EDTA, 25% glycerol, protease inhibitor (Roche)) was then added to the mixture and vortexed vigorously during the 30 min incubation period on ice. The extracted nuclear supernatant was obtained by centrifuging at 21,000 g, 4 °C for 5 min. For SDS-PAGE gel, both the cytosolic pellet and the nuclear pellet were dissolved in 20 μL of 5x SDS gel loading dye and boiled at 95 °C for 5 min. All the supernatant samples (cytosolic, wash, nuclear) were prepared by mixing 20 μL sample with 5 μL 5x SDS gel loading dye and boiled at 95 °C for 5 min. Loading controls were generated by adding 150 nM of purified *t*MeCP2-Strep variants to non-treated nuclear supernatant. Western blot analysis was performed using primary antibodies against Strep-tagII (IBA 2-1509-001), MeCP2 (CST 3456S), LAMP 1 (CST 9091S), EEA1 (CST 3288S), tubulin (CST 2125S), Rab 7 (CST 9367S) and the secondary antibody HRP-linked Anti-rabbit IgG (CST 7074S).

### LC-MS/MS analysis

1 μM ZF-*t*MeCP2-Strep was incubated with Saos-2 cells (37 °C, 5% CO_2_, 1 h) and the nuclear supernatant was isolated as described in the section above. ZF-*t*MeCP2-Strep delivered to the nucleus was enriched by incubating the nuclear supernatant with MagStrep “type3” XT magnetic beads (IBA 2-4090-002) for 45 min at RT in a rotating wheel. After separating the beads from the unbound supernatant using a magnetic separator, the pull-down proteins were eluted with 10 μL 5x SDS-PAGE loading dye (diluted to 2.5x by 10 μL milliQ water) at 95 °C for 3 min. The sample was run on a 10% SDS-PAGE gel (Biorad, Mini-PROTEAN® TGX™) and visualized by Gelcode Blue® Coomassie stain (Pierce) following the manufacturer’s note. The band corresponding to ZF-*t*MeCP2-Strep was extracted from the gel and digested with trypsin, and the resulting peptides were dried and resuspended in buffer: 5% acetonitrile/ 0.02% heptaflurobutyric acid (HBFA)). Mass spectrometry was performed by the Proteomics/Mass Spectrometry Laboratory at UC Berkeley as described in **Supplementary Methods 3**.

### Determining endogenous level of MeCP2 and NCoR

1 × 10^6^ of Saos-2, CHO-K1, NIH3T3 or HeLa cells were lysed with 100 μL M-PER™ Mammalian Protein Extraction Reagent (Thermo Scientific, 78501) for 10 min following the manufacturer’s instructions. The lysates were centrifuged at 14,000 g for 15 min. For SDS-PAGE gel, the pellets were dissolved in 20 μL of 8 M urea and 5 μL 5x SDS gel loading dye and boiled at 95 °C for 5 min. The supernatant samples were prepared by mixing 20 μL sample with 5 μL 5x SDS gel loading dye and boiled at 95 °C for 5 min. Western blot analysis was performed using primary antibodies against MeCP2 (3456S), NCoR1 (CST 5948S), GAPDH (CST 2118S), tubulin (CST 2125S) and the secondary antibody HRP-linked Anti-rabbit IgG (CST 7074S).

### *In cellulo* coimmunoprecipitation assay

On day 1, ∼5 × 10^6^ CHO-K1 cells were plated in 150 mm dishes in F12 Nutrient Mixture (Ham) media with L-Glutamine (+10 % FBS). On day 2, for protein delivery, cells were washed with 25 mL DPBS three times and treated with 1 μM tMeCP2-Strep variants (WT, ZF or Tat, diluted in F12 media without FBS) at 37 °C with 5% CO_2_ for 1 h. Three dishes of cells were incubated with the same F12 media added as controls. After incubation, cells were lifted, washed and lysed as described in the cellular fractionation section. For lysis, the cells from each dish were then suspended in 300 μL of isotonic sucrose buffer (290 mM sucrose, 10 mM imidazole, pH 7.0, 1 mM DTT, and 1 cOmplete protease inhibitor cocktail (Roche)), transferred to 0.5 mL microtubes containing 1.4 mm ceramic beads (Omni International) and homogenized using a Bead Ruptor 4 (Omni International) for 10 s at speed 1. The homogenized cell lysate was centrifuged for 10 min at 800 g, 4 °C. The supernatant contained cytosolic proteins and the crude nuclei pellet was washed in 500 μL of isotonic sucrose buffer supplemented with 0.15% NP-40. After incubating on ice for 10 min, the mixture was centrifuged at 1200 g, 4 °C for 10 min. After separating the wash 1 supernatant, the pellet was washed again with 400 μL of isotonic sucrose buffer supplemented with 0.15% NP-40 and centrifuged at 1200 g, 4 °C for 10 min. The resulting pure nuclear pellet was first resuspended with 70 μL low salt buffer (20 mM HEPES, pH 7.6, 1.5 mM MgCl_2_, 10 mM KCl, 25% glycerol, protease inhibitor (Roche)) and 2.5 μL benzonase (Sigma, 71206), transferred to 0.5 mL microtubes containing 1.4 mm ceramic beads and homogenized using a Bead Ruptor 4 for 10 s at speed 5 for three times (30 s interval on ice between each session). The homogenized nuclear lysate was then incubated at 37 °C for 10 min on a rotating wheel for benzonase digestion. 1M NaCl solution was added to the lysate to a final NaCl concentration of 430 mM. The mixture was homogenized again for 10 s at speed 5 and incubated for 1 h at 4 °C on a rotating wheel. The extracted nuclear supernatant was obtained by centrifuging at 18,000 g, 4 °C for 10 min.

For two control samples, 150 nM purified ZF- or Tat-*t*MeCP2-Strep was doped in nuclear supernatant. The input samples were prepared by taking 12.5 μL of nuclear supernatant and mixing with 7.5 μL milliQ water and 5 μL 5x SDS gel loading dye. The rest of the nuclear supernatant was diluted to 150 mM NaCl with low salt buffer supplemented with 15 mM BME and incubated with MagStrep “type3” XT magnetic beads (IBA 2-4090-002) overnight at 4 °C on a rotating wheel. On the next day, the unbound supernatant was separated from the beads using a magnetic separator. The beads were washed with 200 μL coIP buffer two times (quickly vortex and centrifuge) and eluted with 10 μL 5x SDS-PAGE loading dye (diluted to 2.5x by 10 μL milliQ water) at 95 °C for 3 min. The input and pulldown samples were analyzed by western blot using primary antibodies against Strep-tagII (IBA 2-1509-001), HDAC3 (CST 85057S), TBLR1/TBL1XR1 (CST 74499S), and secondary antibody HRP-linked Anti-rabbit IgG (CST 7074S). To detect for contamination from other cellular compartments, the input and pulldown samples, as well as other fractions during the isolation process (cytosolic supernatant, wash 1, and wash 2), were analyzed by western blot using primary antibodies against EEA1 (CST 3288S), tubulin (CST 2125S), Rab 7 (CST 9367S) and the secondary antibody HRP-linked Anti-rabbit IgG (CST 7074S).

### *In cellulo* transcription assay

To construct the reporter plasmid, the mNeonGreen gene was amplified from a pCS2+mNeonGreen-C Cloning Vector (Addgene #248605) using design primers (**Supplementary Table 3**). The SNRPN promoter was amplified from mouse genomic DNA (Promega, G3091) using design primers (**Supplementary Table 3**). The amplified SNRPN promoter segment was incorporated into a mammalian expression vector pNL1.2 (Promega, N1011) using the two restriction sites NheI and HindIII followed by gibson assembly. The assembled pNL1.2-SNRPN plasmid was confirmed by Sanger sequencing and digested again with HindIII and EcoRI to incorporate the amplified mNeonGreen gene downstream of SNRPN using a gibson assembly kit. The final assembled plasmid was named as pSNRPN-mNeonGreen-PEST. To generate methylated promoter, the plasmid was with HpaII methyltransferase (NEB, M0214S) following manufacturer’s instructions. Complete methylation was checked by testing the plasmid’s resistance to HpaII restriction enzyme digestion.

To perform the assay, on day 1 CHO-K1 cells grown to ∼70% confluency in 60 mm dishes were transfected with either methylated or non-methyalted reporter plasmids in reduced serum media (OPTI-MEM, Gibco, 31985-062) using a TransIT CHO-K1 transfection kit (Mirusb Bio) following manufacturer’s instructions. After 12 h the cells were washed with DPBS two times and lifted with enzyme-free cell dissociation buffer (Gibco). The lifted cells were pelleted, counted and plated into a 24-well plate in 1.2 × 10^5^ cells/well in growth media (F12 Nutrient Mixture (Ham) media (Gibco) with L-Glutamine + 10 % FBS). On day 2, the medium was aspirated and the cells were washed with 1 mL DPBS two times. Then 130 μL of 1-5 μM *t*MeCP2-Strep variants diluted in F12 medium (-FBS, -P/S) were added to the cells and incubated for 1 h at 37 °C, 5 % CO_2_. After delivery, cells were washed with 1 mL of DBPS three times and the media was replaced with the growth media for a one-hour incubation. After incubation, the cells were lifted with 200 μL TrypinLE Express for 5 min at 37 °C, 5% CO_2_, quenched with 1 mL growth media, and transferred to 1.5 mL microcentrifuge tubes. The cells were pelleted by centrifugation at 500 g for 3 min, and washed with 1 mL clear DMEM medium. After centrifuging at 500 g for 3 min, the pellets were resuspended in 200 μL clear DMEM medium and analyzed using an Attune NxT flow cytometer (Life Technologies).

For flow cytometry, the mNeonGreen protein was excited with a laser at 488 nm, and the emission filter was set at 530 ± 30 nm. 130 μL (at least 50,000 cells) were analyzed for each sample and at least three technical and biological replicates were measured for each condition. Cells having green fluorescence were gated based on the background fluorescence level of cells not transfected with reporter plasmids.

### Statistical tests

All statistical tests were done using GraphPad Prism 9 software. Statistical significance for flow cytometry and FCS data were assessed by Brown-Forsythe and Welch ANOVA followed by Dunnett’s T3 multiple comparisons test or unpaired t test with Welch’s correction. The values in the transcriptional repression assay were compared using nonparametric Kruskal-Wallis multiple comparison test. The test used and error bars are defined in each figure legend.

## Data availability

The key datasets generated during and/or analyzed during the current study are included in source data for each figures/tables. Other datasets are available from the corresponding author on reasonable request.

## Code availability

The MATLAB® script used to analyze FCS measurements is available from GitHub (https://github.com/schepartzlab/FCS).

## Acknowledgments

The authors are grateful to all members of the Schepartz lab for helpful discussion. This work was supported by the Rett Syndrome Research Fund. We also thank Dr. Susan Marqusee for the generous use of her CD spectrometer; Anthony Iavarone at QB3/Chemistry Mass Spectrometry Facility at UC Berkeley for mass spectrometry of purified *t*MeCP2 proteins; Dr. Mary West for managing the QB3 Cell and Tissue Analysis Facility at UC Berkeley; Alison Killilea at the UC Berkeley Cell Culture Facility for tissue culture support; Dr. Lori Kohlstaedt at Vincent J. Coates Proteomics/Mass Spectrometry Laboratory at UC Berkeley for LC-MS/MS analysis; Dr. Brian Hodge for training and guidance on RT-qPCR, Jie Fang and Dr. Paul Sauer from Dr. Eva Nogales’s lab for the use of ultracentrifuge; Dr. Bo Li for revising the MATLAB script; Dr. Susan Knox for FCS training and Dr. Sebastian Santiago for protein purification training. Experiment workflow figures were created with Biorender.com.

## Author contributions

A.S. conceived the project. X.Z. designed and conducted experiments. M.Z. performed and interpreted CD analyses. D.M. assisted with the design of *in vitro* pull-down and *in cellulo* co-immunoprecipitation assays. X.Z and A.S. interpreted results, refined experiments, and wrote the manuscript with expert input and editing by M.Z.

## Competing interests

X.Z. and A.S. have filed a provisional patent application related to this work.

**Supplementary information is available for this paper.**

## Materials & Correspondence

Correspondence to Alanna Schepartz.

**Extended Data Fig 1.**
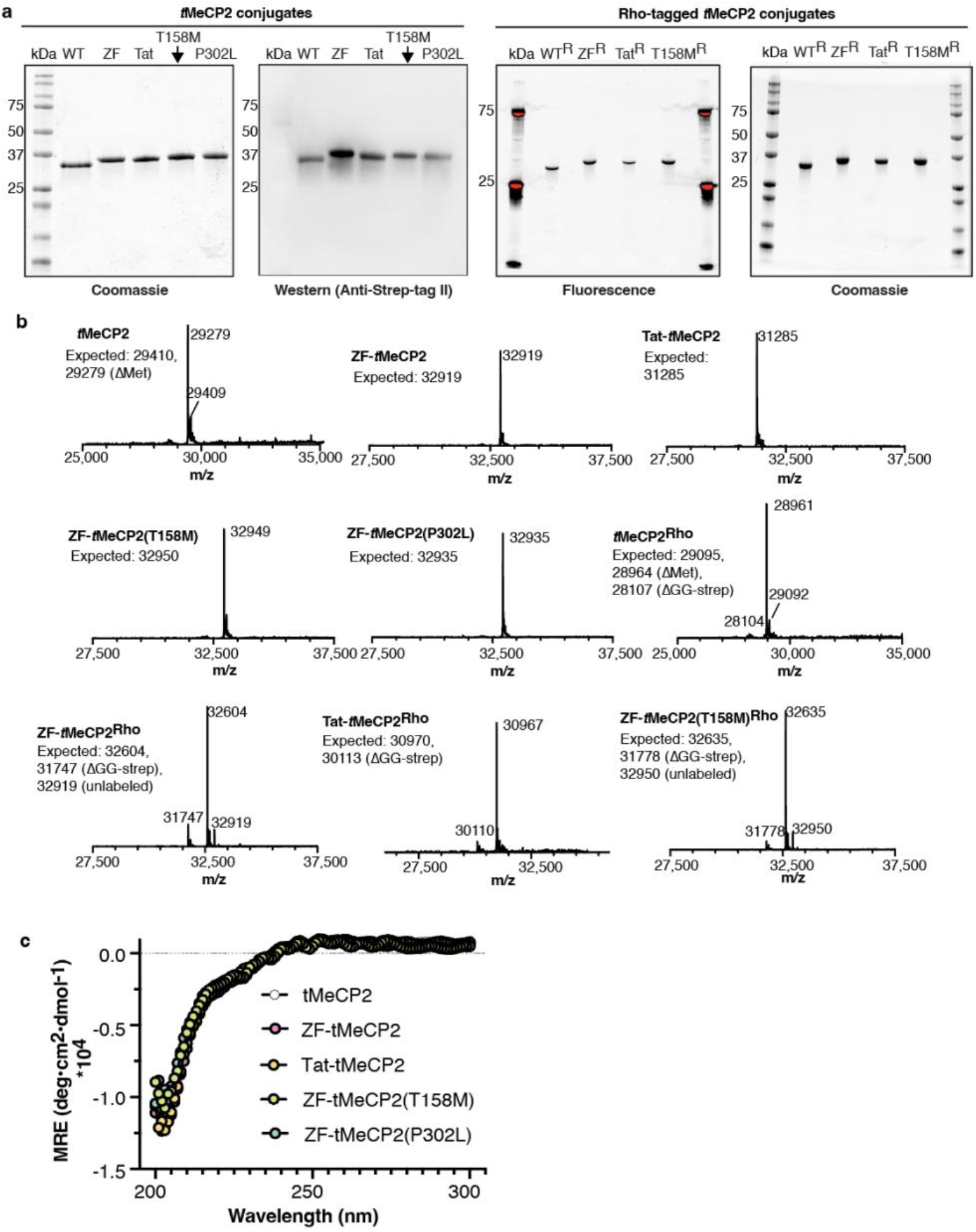
Characterization of purified *t*MeCP2 proteins using SDS-PAGE and mass spectrometry. **a**, (left) Coomassie-stained SDS-PAGE gel and western blot showing the five final purified *t*MeCP2 variants studied herein. (right) Fluorescence imaging confirmed conjugation of Rho to the indicated fusion protein. Labeling efficiencies range from 48-67%. SDS-PAGE shows that purified fusion proteins are all > 90% homogeneous. **b**, LC/MS analysis confirmed the identity of the indicated fusion protein. Prominent peaks were observed for the labeled protein, as well as the presence of unlabeled protein and an intermediate product from sortase labeling. **c**, CD spectra of *t*MeCP2 fusion proteins were collected at ∼12 μM protein concentration in 20 mM HEPES, 300 mM NaCl, 10% glycerol pH 7.6 at RT between 200 and 300 nm at 1 nm intervals with an averaging time of 5 seconds. The protein secondary structure was predicted (**Supplementary Table 1**) using the webserver BeStSel (http://bestsel.elte.hu/index.php).

**Extended Data Fig 2.**
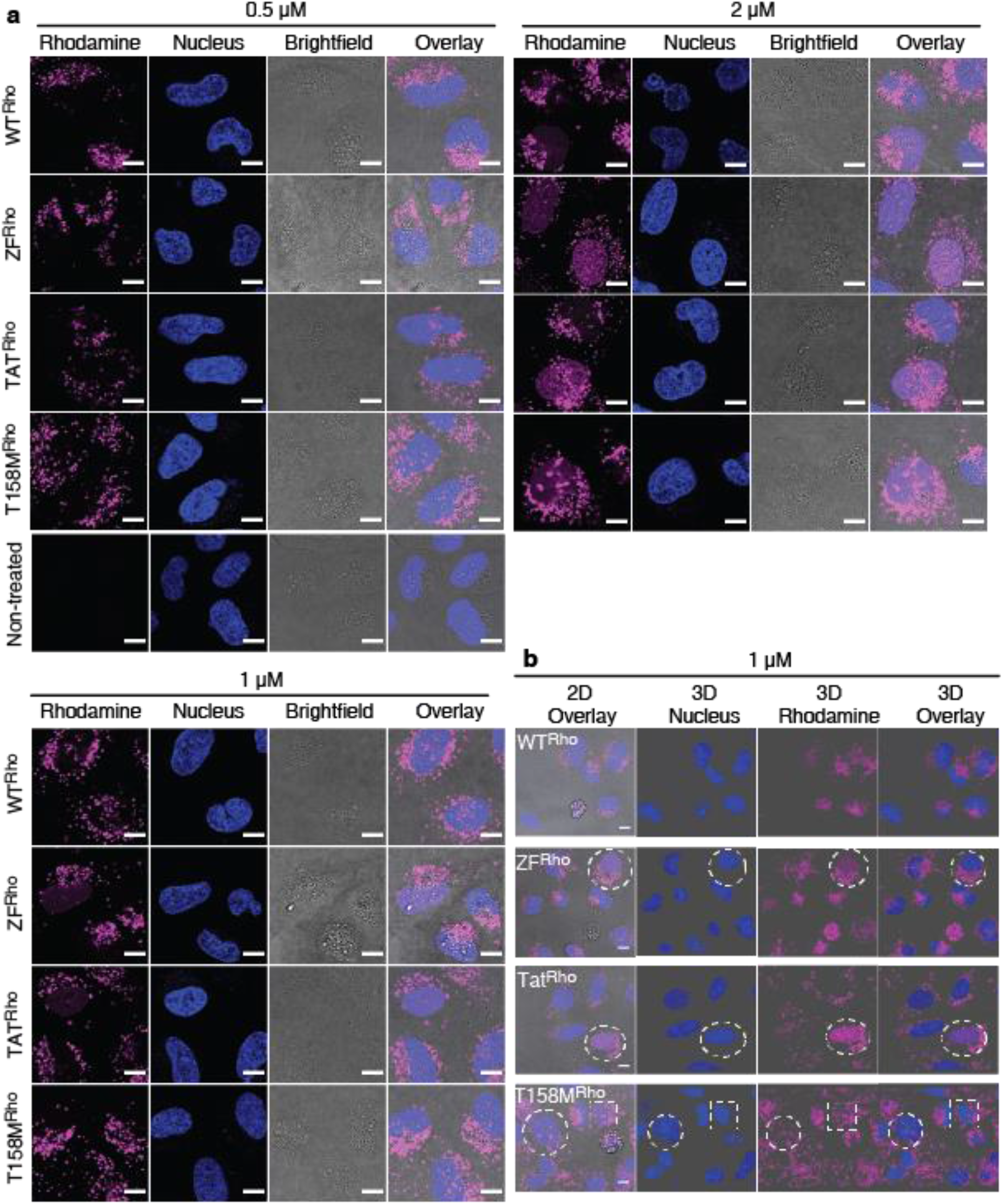
Confocal microscopy images of Saos-2 cells treated with *t*MeCP2-Rho variants. Representative 2D (**a**) and 3D (**b**) confocal images of Saos-2 cells treated with different concentrations of *t*MeCP2-Rho variants (WT^Rho^: *t*MeCP2^Rho^, ZF^Rho^: ZF-*t*MeCP2^Rho^, Tat^Rho^: Tat-*t*MeCP2^Rho^, T158M^Rho^: ZF-*t*MeCP2(T158M)^Rho^) for 1 hr at 37 °C, 5% CO_2_. Nuclei were identified by incubation with Hoechst 33342 for 5 min at the end of the incubation period. Cells with nuclear fluorescence are highlighted in the white dash boxes. Scale bar = 10 μm. The results shown are representative of at least two biological replicates.

**Extended Data Fig 3.**
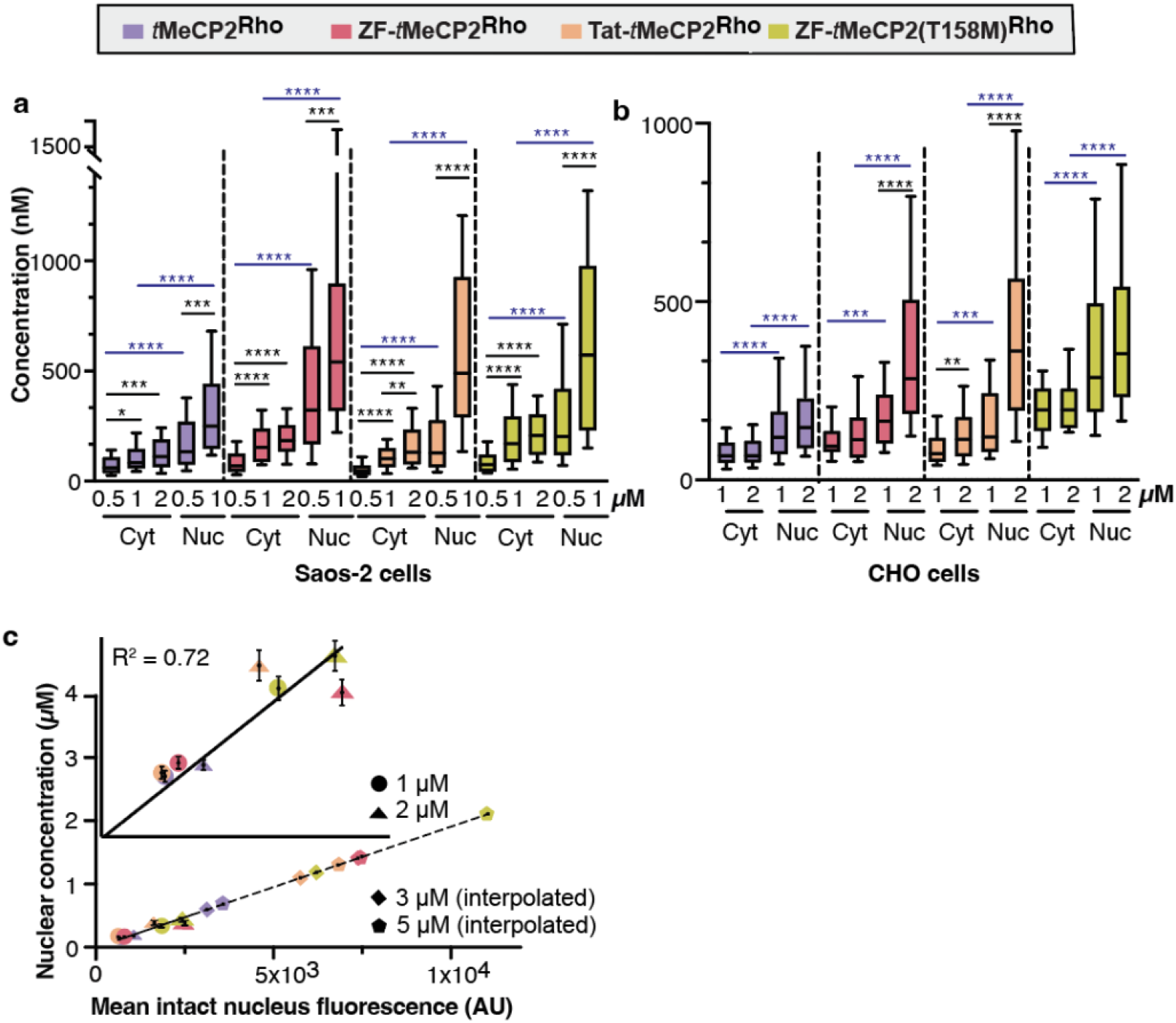
Protein intracellular concentrations determined by FCS. Saos-2 (**a**) or CHO-K1 (**b**) cells were incubated with the indicated *t*MeCP2-Rho variants for 1 hr at 37 °C, 5% CO_2_. The autocorrelation data (**Supplementary Figs. 1-3**) from FCS measurements was fitted to a 3D anomalous diffusion equation (cytosol)^42^ or a two-component 3D diffusion equation (nucleus)^44^ to establish the concentration of each protein in the cytosol and nucleus. Center line, median; box limits, 25-75 percentile; whiskers, 10-90 percentile. (n > 30 cells total for each condition from at least two biological replicates, see source data). The cytosolic and nuclear concentrations of the same *t*MeCP2-Rho variant under the same treatment condition (e.g. 1 μM) were compared using two tailed unpaired parametric t test with Welch’s correction. The dose-dependent concentrations of the same *t*MeCP2-Rho variant under the same location (e.g. WT^Rho^ in the cytosol) were compared using either Brown-Forsythe and Welch ANOVA followed by Dunnett’s T3 multiple comparisons test (for cytosolic concentration in Saos-2 cells, P values are adjusted to account for multiple comparisons) or two tailed unpaired parametric t test with Welch’s correction (for nuclear concentration in Saos-2 cells and all CHO-K1 cell data) ****p ≤ 0.0001, ***p ≤ 0.001,**p ≤ 0.01, *p ≤ 0.05. (see source data). **c**, Scatter plot depicting the correlation of the mean nuclear fluorescence of intact CHO-K1 nuclei measured by flow cytometry (x-axis) and the intra-nuclear concentrations measured by FCS (y-axis). Dots are represented as mean ± SEM. Inset: Data for treatment with 1-2 μM proteins were fitted with a simple linear regression line. The nuclear concentrations of CHO-K1 cells treated with 3 μM and 5 μM proteins (diamonds, pentagons) were interpolated based on their corresponding nuclear fluorescence values.

**Extended Data Fig. 4.**
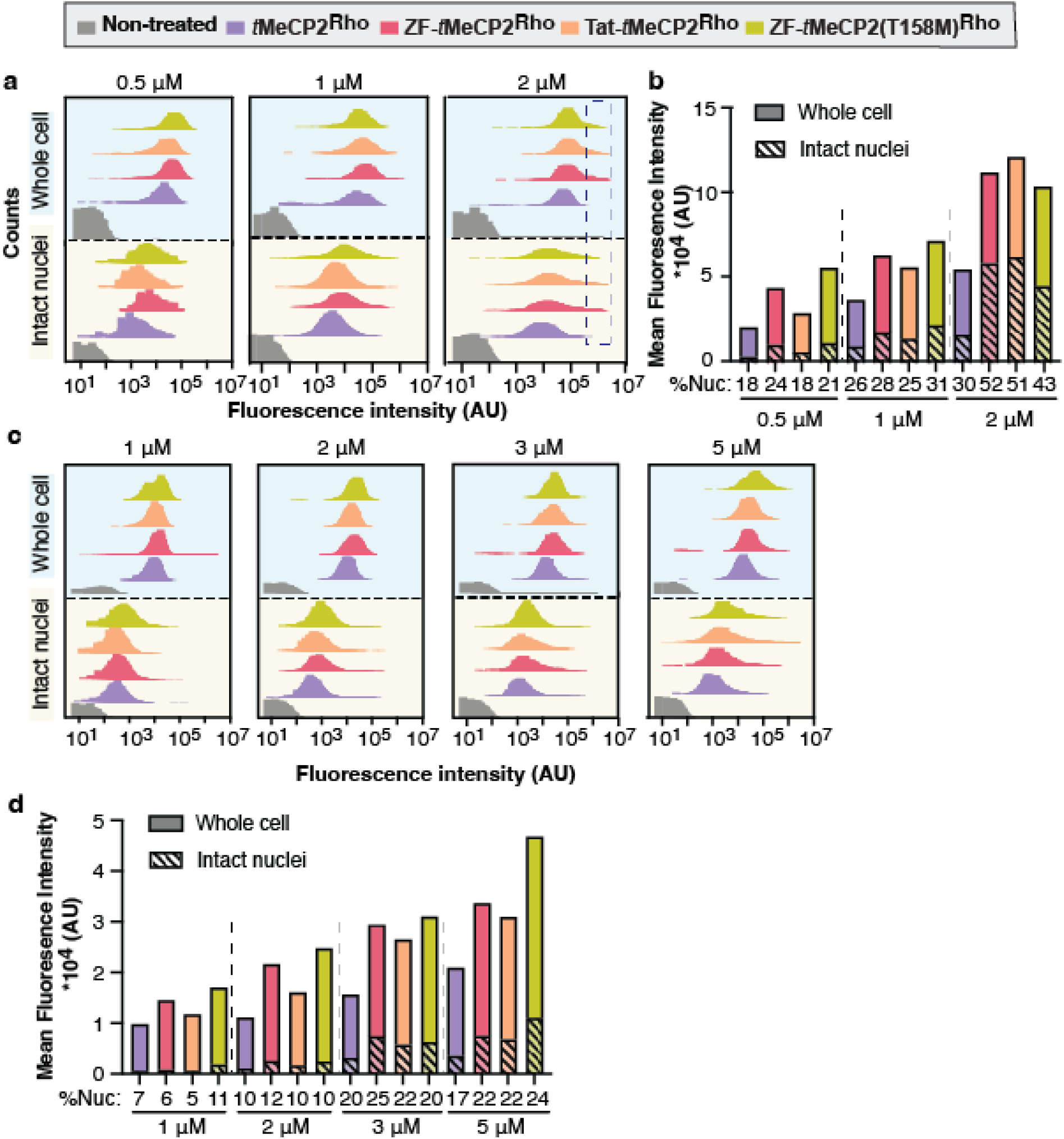
Flow cytometry analysis of Saos-2 and CHO-K1 cells treated with MeCP2-Rho variants. Whole cells and intact nucleus rhodamine fluorescence histogram of Saos-2 cells (**a**) and CHO-K1 cells (**c**) treated with MeCP2-Rho variants for 1 hr at 37 °C, 5% CO_2_ measured by flow cytometry, with the mean rhodamine fluorescence intensity and percentage of nuclear delivery shown in (**b**) and (**d**). n>9000 and n>1500 for whole cell and intact nuclei flow cytometry, respectively, see source data.

**Extended Data Fig 5.**
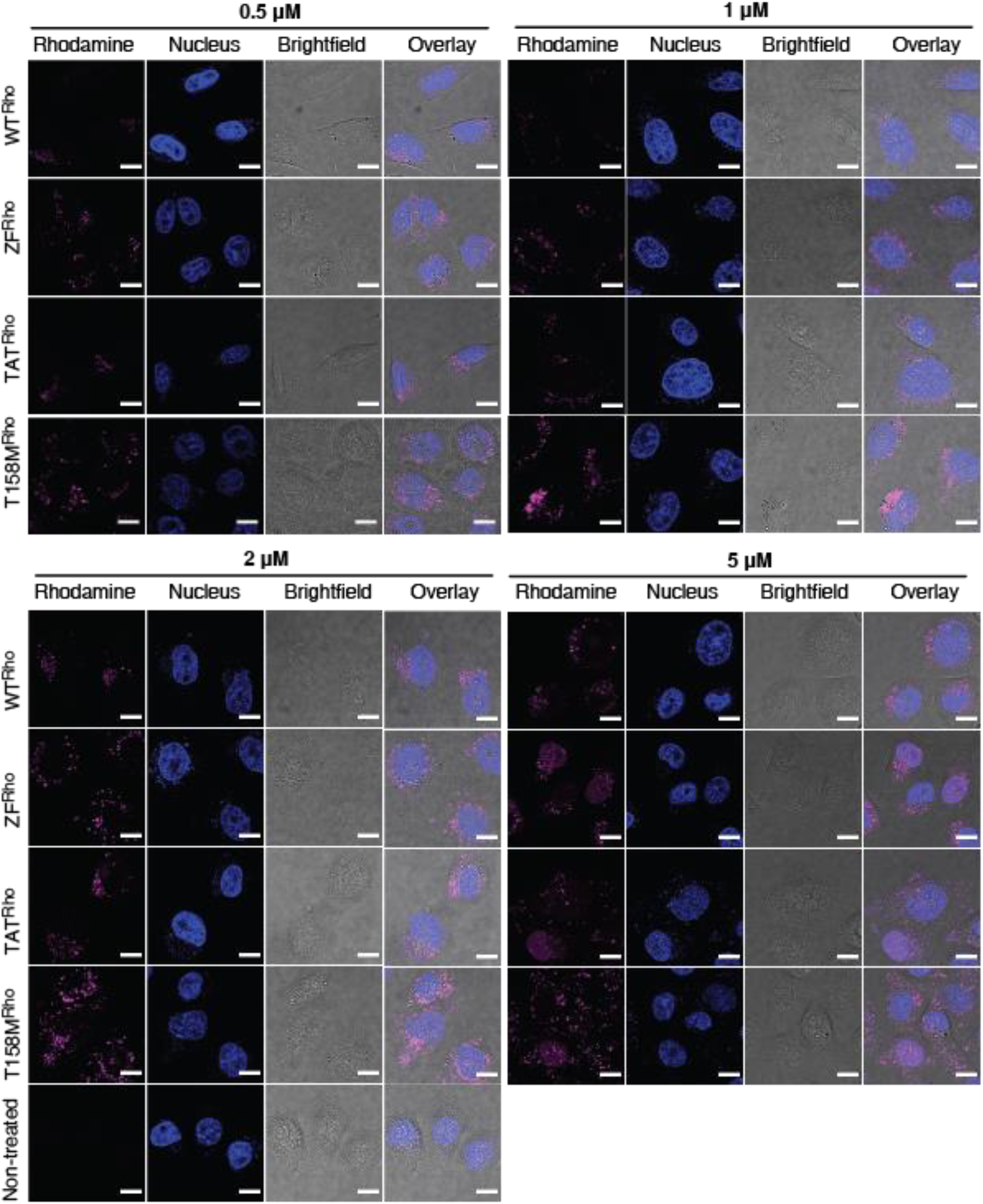
Representative 2D confocal images of CHO-K1 cells treated with *t*MeCP2-Rho variants. Different concentrations of *t*MeCP2-Rho variants (WT^Rho^: *t*MeCP2^Rho^, ZF^Rho^: ZF-*t*MeCP2^Rho^, Tat^Rho^: Tat-*t*MeCP2^Rho^, T158M^Rho^: ZF-*t*MeCP2(T158M)^Rho^)) were added to the cells and incubated for 1 hr at 37 °C, 5% CO_2_. Nuclei were identified by incubation with Hoechst 33342 for 5 min at the end of the incubation period. Scale bar = 10 μm. The results shown are representative of at least two biological repeats.

**Extended Data Fig 6.**
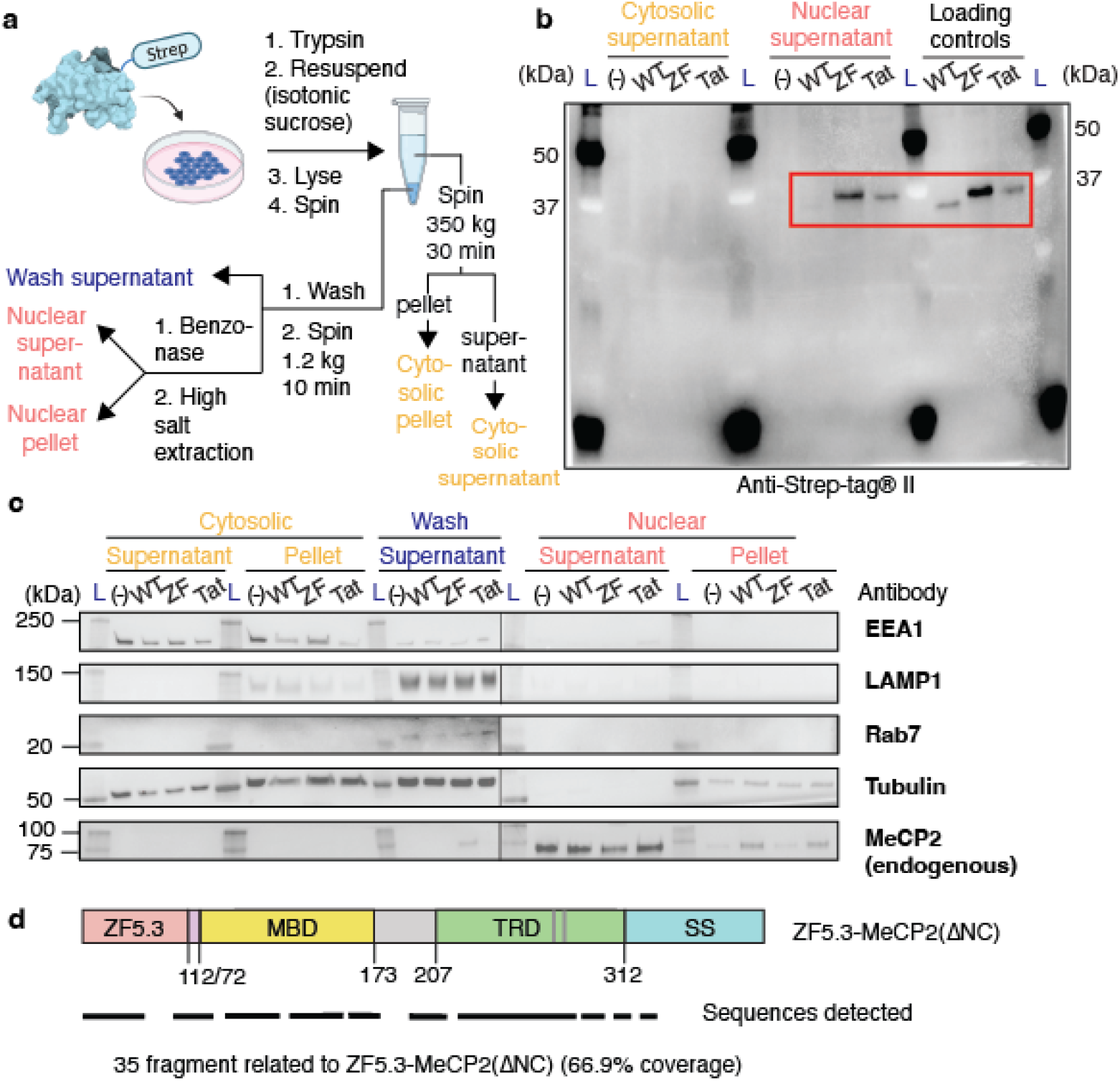
Cellular fractionation and LC-MS/MS analysis of delivered intact proteins. (**a**) Scheme illustrating the workflow to isolate the cytosolic and nuclear fractions of Saos-2 cells after delivery of 1 μM *t*MeCP2 (WT), ZF-*t*MeCP2 (ZF), Tat-*t*MeCP2 (Tat) for 1 hr at 37 °C, 5% CO_2_. One dish of cells with the same clear McCoy’s media added was included as a non-treated control (-). Detailed isolation procedures can be found in Methods and Materials. (**b**) Western blot analysis of the cytosolic and nuclear supernatants isolated from (**a**) using an antibody against Strep-tagII (IBA 2-1509-001). For loading controls, the nuclear supernatant of non-treated cells were doped with 150 nM of purified *t*MeCP2, ZF-*t*MeCP2, and Tat-*t*MeCP2. Bands corresponding to intact ZF-*t*MeCP2 and Tat-*t*MeCP2 in both the nuclear supernatant and loading control are highlighted in the red box. L, Ladder. (**c**) Western blot analysis of different cellular fractions isolated from (**a**) with antibodies against endocytic (EEA1, LAMP1, Rab7), cytosolic (tubulin) and nuclear (endogenous MeCP2) markers. The gel results shown are representative of three biological repeats. L, Ladder. (**d**) Scheme illustrating the positions of fragments detected by LC-MS/MS. ZF-tMeCP2 isolated from Saos-2 cells as shown in (**a**) was enriched with MagStrep “type3” XT magnetic beads (IBA 2-4090-002) that recognizes the C terminal Strep-tagII and subjected to trypsin digest before LC-MS/MS analysis. In total 35 relevant fragments were identified, corresponding to 66.9% of protein coverage. Representative spectra are shown in **Supplementary Figure 5**.

**Extended Data Fig 7.**
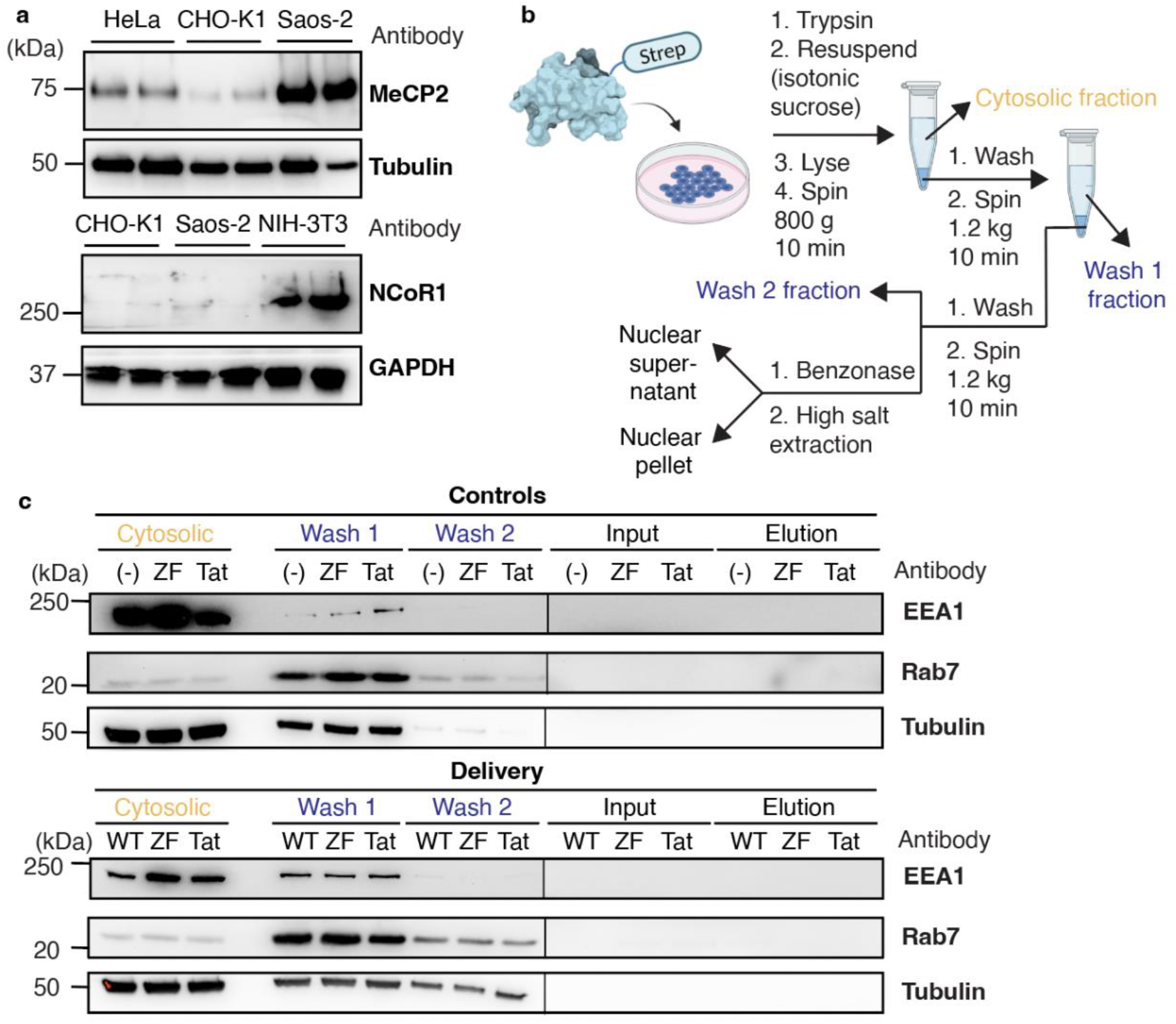
*In cellulo* co-immunoprecipitation assay in CHO-K1 cells. **a**, Relative levels of endogenous MeCP2 and NCoR1 in lysates prepared from different cell lines. Western blots were performed using primary antibodies against MeCP2 (3456S), tubulin (CST 2125S), NCoR1 (CST 5948S), GAPDH (2118S) and the secondary antibody HRP-linked Anti-rabbit IgG (CST 7074S). **b**, Scheme illustrating the workflow to isolate and extract the nuclear proteins of CHO-K1 cells after incubation with 1 μM *t*MeCP2 (WT), ZF-*t*MeCP2 (ZF), Tat-*t*MeCP2 (Tat) for 1 hr at 37 °C, 5% CO_2_. Three dishes of non-treated cells (-) were incubated under identical conditions as controls and later doped with either 150 nM ZF-*t*MeCP2 or Tat-*t*MeCP2 after the nuclear supernatant fractions were isolated. Detailed isolation procedures can be found in Methods and Materials. **c**, Western blot analysis of different cellular fractions isolated from (**b**) with antibodies against endocytic (EEA1, Rab7) and cytosolic (tubulin) markers. The gel results shown are representative of two biological repeats.

**Extended Data Fig 8.**
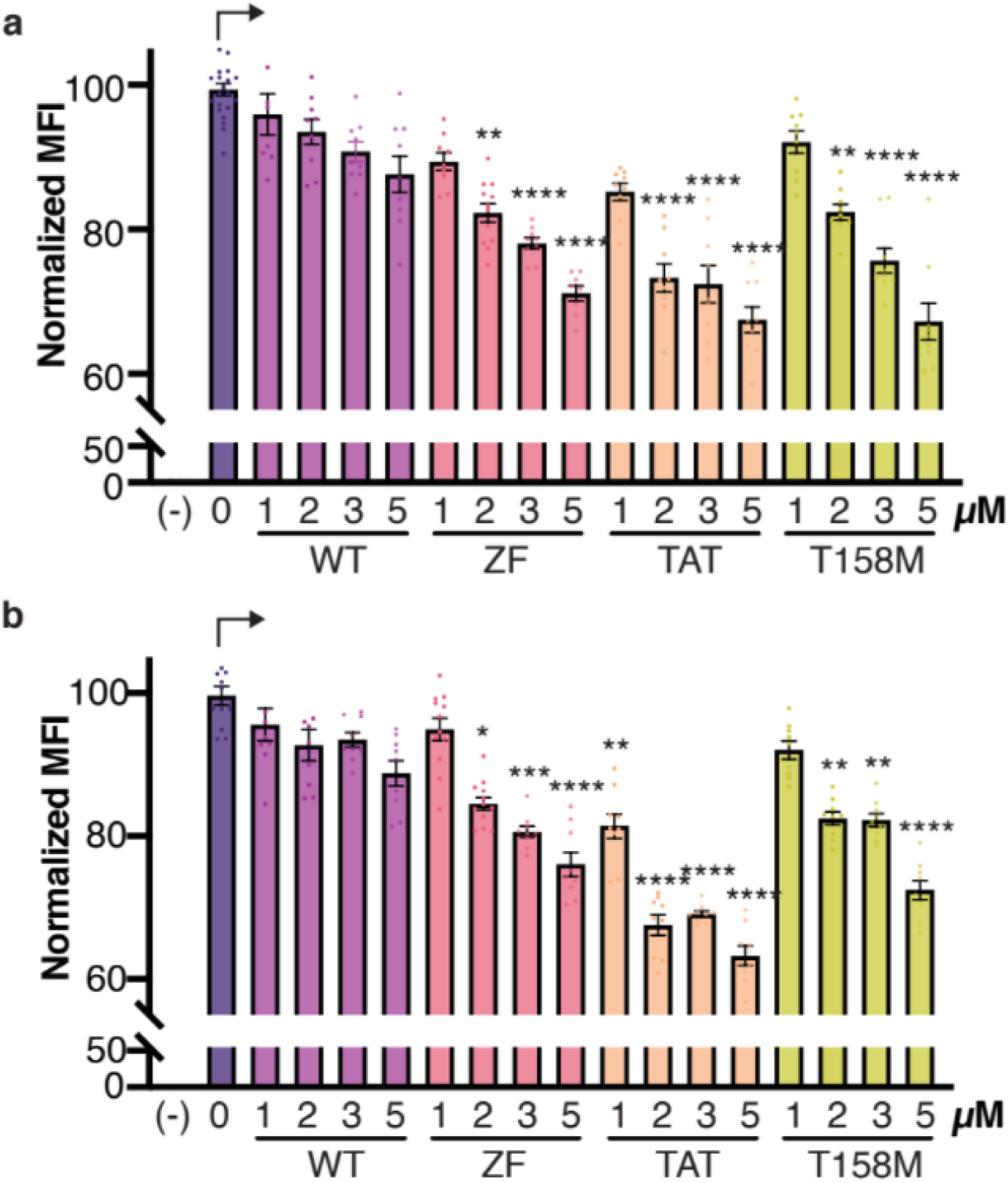
Mean fluorescence intensities (MFI) of CHO-K1 cells studied in transcription repression assay. CHO-K1 cells transfected with methylated (**a**) or non-methylated **(b)** reporter plasmids were incubated with 1-5 μM the indicated tMeCP2 variants (WT: tMeCP2, ZF: ZF-tMeCP2, Tat: Tat-tMeCP2, P: ZF-tMeCP2(P302L)) for 1 hr at 37 °C, 5% CO_2_ and analyzed using flow cytometry to identify changes in green fluorescence levels. Cells were gated for green fluorescence based on the background fluorescence level of cells that were not transfected with a reporter plasmid. Data are represented as mean ± SEM. Each sample comprised 130 μL (at least 50,000 cells) and at least three technical and biological replicates at each condition, see source data. The MFI was normalized against non-treated cells transfected with the reporter plasmid. Statistical significance compared with non-treated cells transfected with the reporter plasmid was calculated using a nonparametric Kruskal-Wallis test followed by Dunnett’s T3 multiple comparisons test. ****p ≤ 0.0001, ***p ≤ 0.001,**p ≤ 0.01, *p ≤ 0.05. Each P value is adjusted to account for multiple comparisons (see source data).

**Extended Data Table 1.** Intracellular concentrations calculated from FCS measurements. Saos-2 or CHO-K1 cells were incubated with the indicated *t*MeCP2-Rho variants for 1 hr at 37 °C, 5% CO_2_. The autocorrelation data (**Supplementary Figs. 1-3**) from FCS measurements was fitted to a 3D anomalous diffusion equation (cytosol)^1^ or a two-component 3D diffusion equation (nucleus)^2^ to establish the concentration of each protein in the cytosol and nucleus.

| Saos-2 cells |  |  |  |  |  |  |  |  |
| --- | --- | --- | --- | --- | --- | --- | --- | --- |
|  | WT-Cyt | WT-Nuc | ZF-Cyt | ZF-Nuc | Tat-Cyt | Tat-Nuc | T158M-Cyt | T158M-Nuc |
| 0.5 μM | | | | | | | | |
| Number of values | 52 | 72 | 61 | 63 | 51 | 57 | 60 | 73 |
| Median | 60.7 | 134 | 68.5 | 322 | 46.8 | 128 | 74.5 | 202 |
| Mean | 74.1 | 183 | 85.1 | 422 | 52.9 | 183 | 87.8 | 289 |
| Std. Error of Mean | 6.82 | 16.5 | 7.26 | 42.6 | 4.82 | 20.6 | 7.2 | 27.3 |
| 1 μM | | | | | | | | |
| Number of values | 52 | 52 | 47 | 60 | 47 | 58 | 95 | 110 |
| Median | 83.1 | 250 | 153 | 541 | 102 | 490 | 170 | 572 |
| Mean | 107 | 314 | 174 | 709 | 109 | 625 | 208 | 670 |
| Std. Error of Mean | 9.03 | 28.1 | 15.5 | 68.5 | 9.53 | 62.6 | 15.4 | 47.4 |
| 2 μM | | | | | | | | |
| Number of values | 44 | 45 | 29 |  | 84 |  | 65 |  |
| Median | 111 | 342 | 184 |  | 131 |  | 207 |  |
| Mean | 132 | 552 | 198 |  | 161 |  | 225 |  |
| Std. Error of Mean | 12.7 | 64.2 | 16 |  | 11.3 |  | 15.5 |  |
| CHO-K1 cells |  |  |  |  |  |  |  |  |
|  | WT-Cyt | WT-Nuc | ZF-Cyt | ZF-Nuc | Tat-Cyt | Tat-Nuc | T158M-Cyt | T158M-Nuc |
| 1 μM | | | | | | | | |
| Number of values | 66 | 64 | 47 | 52 | 39 | 45 | 49 | 72 |
| Median | 67.94 | 119.5 | 94.93 | 164.4 | 74.1 | 120.7 | 196 | 287.2 |
| Mean | 79.69 | 151.9 | 110.2 | 186 | 90.02 | 160.7 | 205.5 | 372.8 |
| Std. Error of Mean | 5.624 | 14.01 | 7.707 | 15.53 | 7.661 | 15.69 | 14.76 | 29.86 |
| 2 μM | | | | | | | | |
| Number of values | 53 | 81 | 49 | 64 | 54 | 69 | 56 | 72 |
| Median | 68.2 | 147.3 | 112 | 284.4 | 113.7 | 361.3 | 196.6 | 353.7 |
| Mean | 83.48 | 180.2 | 132.5 | 362.2 | 129.1 | 430.2 | 214.5 | 454.5 |
| Std. Error of Mean | 6.386 | 12.76 | 12.52 | 32.54 | 10.58 | 38.16 | 11.46 | 37.95 |

**Extended Data Table 2.**
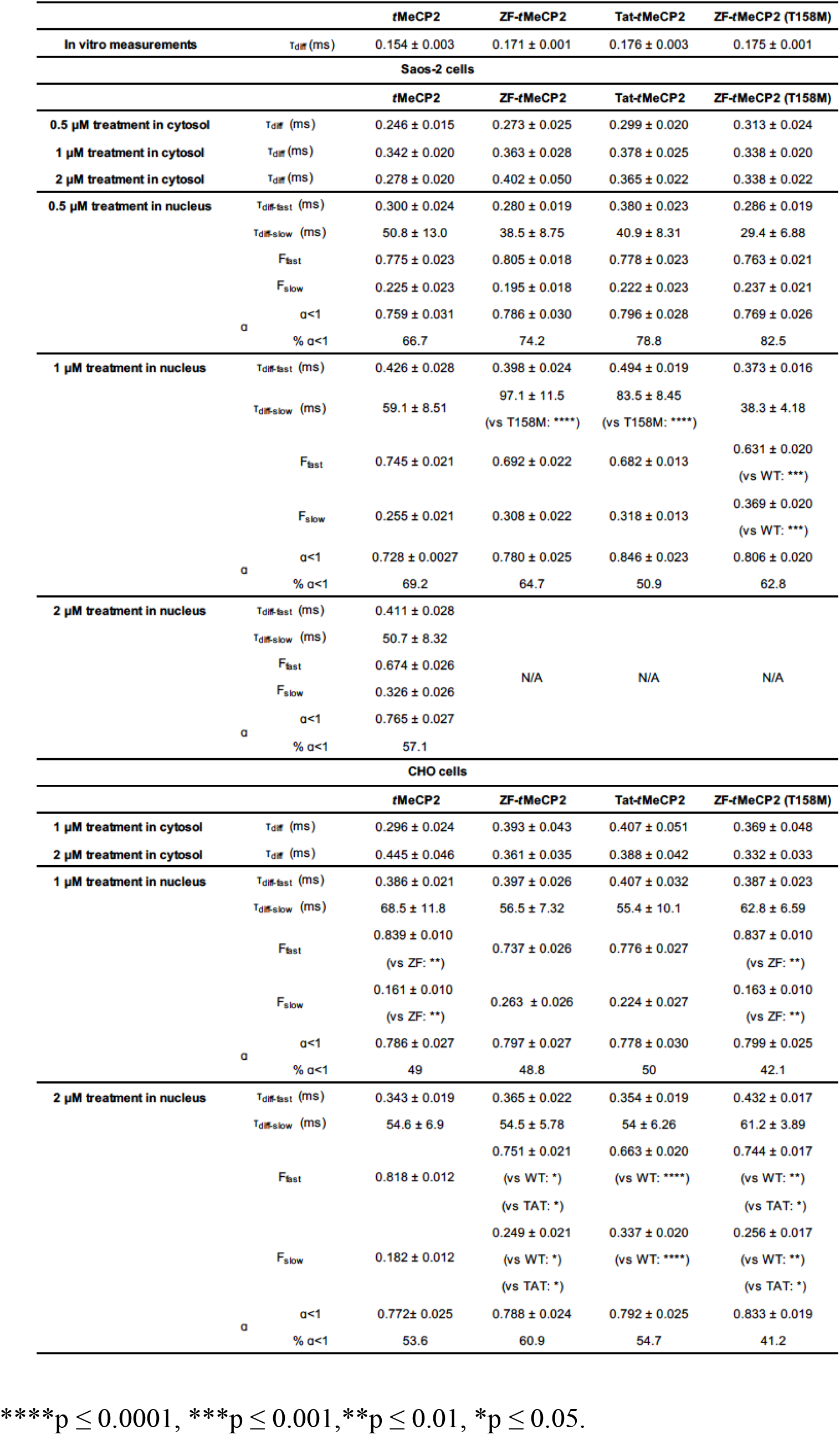
Fitting parameters for FCS measurements. The autocorrelation curves obtained from the *in vitro* measurements were fitted to a 3D diffusion equation and the *in vivo* measurements were fitted to a 3D anomalous diffusion equation (cytosol) and a two-component 3D diffusion equation (nucleus). Values are represented in mean ± SEM. (n>20 from at least two biological replicates, see source data). The values of T_diff-slow_, F_fast_ and F_slow_ in each condition are compared using Brown-Forsythe and Welch ANOVA followed by Dunnett’s T3 multiple comparisons test.

